# Microplastic exposure alters immune-related gene expression in *Culex quinquefasciatus* mosquitoes and larval microbiota diversity

**DOI:** 10.64898/2026.08.27.747521

**Authors:** Ahamed Tchatakoura, Marie Buysse, Marie-Laure Setier-Rio, Olivier Roux, Claire Loiseau, Amandine Aviles

**Affiliations:** UMR MIVEGEC, Université de Montpellier, IRD, CNRS – Montpellier, France; Entente Interdépartementale pour la démoustication du littoral méditerranéen – Montpellier, France

**Keywords:** Culex quinquefasciatus, polyethylene, RNA sequencing, microbiota

## Abstract

**Background:** Microplastics have been detected in many freshwater ecosystems, including stagnant waters where mosquito larvae develop. Larvae are therefore exposed to microplastic pollution, which may affect their life history traits or microbiota along with immune gene expression. However, these effects have never been tested in mosquitoes, despite their major public health importance as vectors of numerous pathogens.

**Method:** We exposed mosquito larvae, from hatching to adult emergence, to four concentrations of polyethylene microplastics (MPs): 0, 60, 200, and 600 MPs/mL. Fourth-instar larvae and newly emerged adult females were collected for each treatment. To investigate the effects of MPs on gene expression and bacterial microbiota, RNA sequencing and 16S metabarcoding approaches were performed on three biological replicates for each developmental stage.

**Results:** Microplastic exposure induced a non-monotonic dose-dependent transcriptomic response. In larvae, only a limited number of genes were differentially expressed (five to eight per concentration), with immune-related genes downregulated at both low and high concentrations. In adult females, the intermediate concentration (200 MPs/mL) elicited the strongest response with 14 differentially expressed genes (DEGs). Regarding the microbiota, microplastic exposure reduced bacterial diversity in larvae, with the lowest diversity observed at the highest concentration. However, no significant changes were detected in the microbiota of adult females.

**Conclusion:** Overall, this study shows that adult females are affected by larval exposure to MPs (*i.e.* differential expression in immune-related genes) and warrants further studies in this field. This includes: 1) investigating further the effects of MPs on mosquitoes’ populations (e.g. through multi-generational studies), 2) gaining more environmentally relevant knowledge on MP effects (*i.e.* using MPs with a biofilm and/or adsorbed pollutants) and 3) focusing on the effects of MPs on mosquitoes’ vectorial capacities.

## Introduction

Over the past decades, plastics have become among the most extensively used materials worldwide. Global annual production doubled between 2000 and 2019, increasing from 234 to 460 million tons (Mt) **(OECD, 2022a)**. Projections indicate that plastic use is expected to nearly triple, rising from 460 Mt in 2019 to 1,231 Mt by 2060 **(OECD, 2022b)**. This massive use of plastic products has inevitably resulted in equally massive waste: an estimated 22% of plastics end up in the environment annually **(OECD, 2022a)**. Once discarded, plastics gradually degrade into microplastics (particles smaller than 5 mm) and nanoplastics (smaller than 1 μm), which infiltrate ecosystems and food chains, affecting organisms ranging from zooplankton to vertebrates (Arif et al., 2024).

The ubiquitous presence of microplastics (MPs) in marine and freshwater environments promotes their interaction with a wide range of organisms, which can ingest them **(Lu et al., 2018; Wang et al., 2021)**. Such ingestion has been associated with various detrimental effects **(Franzellitti et al., 2019),** including inflammation **(Limonta et al., 2019)**, tissue alterations **(Nguyen et al., 2026)**, and physiological impairment in several species **(Banaee et al., 2025)**. Chronic exposure has even been linked to a fibrotic disease termed ‘plasticosis’ in seabirds **(Arif et al., 2024; Charlton-Howard et al., 2023; Lee et al., 2025)**. Beyond direct ingestion, MPs may exert indirect effects through the transport of toxic compounds, such as pesticides (**Sahai et al., 2023**) or persistent organic pollutants (**Koelmans et al., 2013**) and microorganisms (i.e., the plastisphere **Shi et al., 2023**), thereby disrupting physiological functions **(Aryaprema et al., 2025; Rafa et al., 2024)**.

Among freshwater organisms exposed to MPs, mosquitoes occupy a key position because of their major role in public health. Three mosquito genera in particular (*Aedes*, *Culex*, and *Anopheles*) are vectors of numerous pathogens, such as arboviruses (e.g. dengue, chikungunya and West Nile viruses) as well as malaria parasites. Depending on the species, eggs are laid either on a moist substrate or on the water surface. Larvae and pupae develop in aquatic environments prior to adult emergence. MPs have been detected in all habitats colonized by mosquito larvae, ranging from stagnant freshwater and brackish environments (e.g. ponds, ditches, marshes) to highly anthropized habitats such as storm drains, water-filled containers and used tires **(McConnel et al., 2024)**. Morever, female mosquitoes do not avoid water contaminated with MPs when ovipositing **(Cuthbert et al., 2019)** and, while mosquito larvae primarily feed on microorganisms, organic debris, and algae (**Souza et al. 2019**), they can also ingest MPs present in their environment when they are sufficiently small **(Yee et al., 2004)**.

Studies have shown that MPs ingested by mosquito larvae can accumulate in the midgut and Malpighian tubules and be transferred ontogenetically from the larval to the adult stage **(Al-Jaibachi et al., 2018, 2019)**. A limited number of studies have examined the effects of MPs on mosquito’s life-history traits, reporting either no effects on development or survival **(Thormeyer & Tseng, 2023)** or reductions in developmental rate, emergence success or fecundity **(Aryaprema et al., 2025; Li et al., 2024)**. **Edwards et al. (2023)** showed that MP ingestion by *Aedes aegypti* and *Aedes albopictus* larvae significantly disrupts their microbiota, decreasing the abundance of certain commensal bacteria (*e.g. Wolbachia*, *Filimonas*) while increasing the relative abundance of genera such as *Serratia* or *Elizabethkingia*. Similarly, **Li et al. (2024)** observed that MPs ingestion alters both bacterial composition and gut bacteria diversity in *Cx. quinquefasciatus*.

Such microbiota changes could have major repercussions on vectorial capacity, defined as the potential of a mosquito population to transmit pathogens **(Cansado-Utrilla et al., 2021)**. In vector mosquitoes, the microbiota plays a central role in host physiology, development and immunity **(Gao et al., 2020)**. Bacteria can influence pathogen replication within the host and their transmission. For example, *Wolbachia pipientis* and *Chromobacterium sp.* can inhibit viral replication, whereas other bacteria (*e.g. Serratia marcescens*) can promote arbovirus infection **(Apte-Deshpande et al., 2014; Gabrieli et al., 2021; Wu et al., 2019)**. Similarly, *Elizabethkingia anophelis* can influence both *Plasmodium* development in *Anopheles* mosquitoes and Zika virus infection in *Ae. albopictus* **(Bahia et al., 2014; Onyango et al., 2021)**. These effects arise from complex interactions among the microbiota, pathogens and the host innate immune system, orchestrated through two major signaling pathways: the Toll pathway, which induces antimicrobial peptides (AMPs) such as *Cecropins* and *Defensins,* and the Immune Deficiency pathway (IMD), which regulates peptides including *Attacins* and *Gambicins* **(Gabrieli et al., 2021)**. These pathways act directly against bacteria, fungi and certain parasites **(Hillyer, 2010; Kumar et al., 2018; Lv et al., 2023)**, but are also activated by environmental abiotic stresses, thereby modulating the expression of genes involved in immune responses and detoxification **(Reid et al., 2018)**. For example, exposure to insecticides or temperature variation can trigger molecular responses that alter microbiota composition and modulate host gene expression **(Lv et al., 2023; Reid et al., 2018)**.

Although microplastic-induced changes in gene expression have been documented in freshwater insects (e.g. chironomids: **Carrasco-Navarro et al., 2021; Doria et al., 2025**), integrative studies combining transcriptomics and microbiota analyses remain scarce. Only one recent study reported that polystyrene (PS) MPs triggered biochemical responses in damselfly larvae, *Ischnura elegans*, increasing oxidative stress and altering amino acid and fatty acid metabolism pathways. PS exposure was found to modify also the gut microbiota of these larvae, with increased abundances of Firmicutes and Proteobacteria (L. Sun et al., 2025). To date, no comparable studies have been conducted in mosquitoes’ larvae. In this context, our study aims to explore the interactions between microplastic ingestion, bacterial communities and gene expression in both larvae and adult females of *Culex quinquefasciatus*, a mosquito species that vectors several pathogens including West Nile virus (WNV) and *Plasmodium relictum*, the causative agent of avian malaria. We combined RNA-seq transcriptomics and 16S rRNA metabarcoding to characterize MP-induced changes across different MPs concentrations in larvae and adults, with a particular focus on (i) the expression of immunity-related genes and (ii) bacterial diversity community composition. Based on the literature, we hypothesize that larval exposure to MPs could disrupt both immune and detoxification-related gene expression, as well as gut microbiota composition, which could ultimately affect vectorial capacity **(Loiseau & Sorci, 2022)**. Specifically, we anticipate that MPs exposure may alter the abundance of key bacterial taxa such as *Wolbachia* and *Elizabethkingia* **(Edwards et al., 2023; Li et al., 2024)** while modulating the expression of immune-related genes and stress-response pathways.

## Material and methods

### Mosquito strain and rearing conditions

A laboratory strain of *Cx. quinquefasciatus* Slab, provided by the Entente Interdépartementale pour la Démoustication du littoral Méditerranéen (EID-Med), was used in this study. At EID-Med, mosquitoes are reared under controlled conditions at a constant temperature 27 ± 1°C and 70 ± 10% relative humidity and a 16:8 h (light:dark) photoperiod. Larvae were fed *ad libitum* with guinea pig pellets (Safe – Premium Scientific Diet). Adult mosquitoes were provided with honey water and adult females received blood meals using an artificial blood-feeding device. All rearing equipment used for this study was made of glass. Eggs were collected in glass containers, and first-instar larvae were used for exposure immediately after hatching.

### Microplastic preparation

Polyethylene (PE) MPs (30-50 µm, Sigma-Aldrich, CAS 9002-88-4, density 0.94 g/mL) were used to establish four experimental conditions. MPs were suspended in 0.1X PBS to prepare a stock solution at a concentration of 100,000 MPs/mL, from which three exposure concentrations (60, 200, and 600 MPs/mL) were prepared. The control consisted of glass beakers with no MPs (0 MPs/mL) but a volume of 0.1X PBS corresponding to the highest volume used to prepare the beakers with MPs (**Figure S1**). Polyethylene was selected because it is one of the most prevalent plastic polymers found in aquatic environments **(Rodrigues Dos Santos et al., 2023)**. The concentrations tested fall within the range reported in the literature for experimental studies

**(Edwards et al., 2023; Griffin et al., 2023; Li et al., 2024)** while also encompassing environmentally relevant concentrations reported in urban habitats. For instance, concentrations ranging from 22 MPs/mL to 2,580 MPs/mL, with an average of 493 MPs/mL, have been reported in storm drains **(Iannuzzi, 2025)**.

### MP exposure experiment

All exposure experiments were conducted in glass beakers of 250mL, filled with 200mL of reverse osmosis water, to minimize potential plastic contamination (**Figure S1**). Beakers were maintained at 26 ± 1°C, 70 ± 10% relative humidity and a 16:8 h (light:dark) photoperiod, corresponding to the rearing conditions. Larvae were fed *ad libitum* with guinea pig pellets (Safe – Premium Scientific Diet) during the experiment.

For each MP concentration condition, seven to nine beakers, each containing 20 larvae, were monitored from hatching to adult emergence to ensure sufficient biological replication for downstream analyses. Three biological replicates per treatment were collected at the fourth-instar larval stage (L4, pre-pupation), with each replicate consisting of a pool of three larvae originating from the same beaker. Similarly, three biological replicates per treatment were collected for newly emerged adult females; each replicate consisting of a pool of three females from the same beaker.

Larvae and female mosquitoes were euthanized at –20°C and immediately preserved in DNA/RNA shield buffer (Zymo Research) at –20°C during sample collection, before being transferred to –80°C for storage until DNA (16S rRNA metabarcoding) and RNA (RNA-seq) extractions (**Figure 1**). Remaining females from each treatment were dissected to isolate midguts for 16S rRNA metabarcoding.

**Figure 1.**
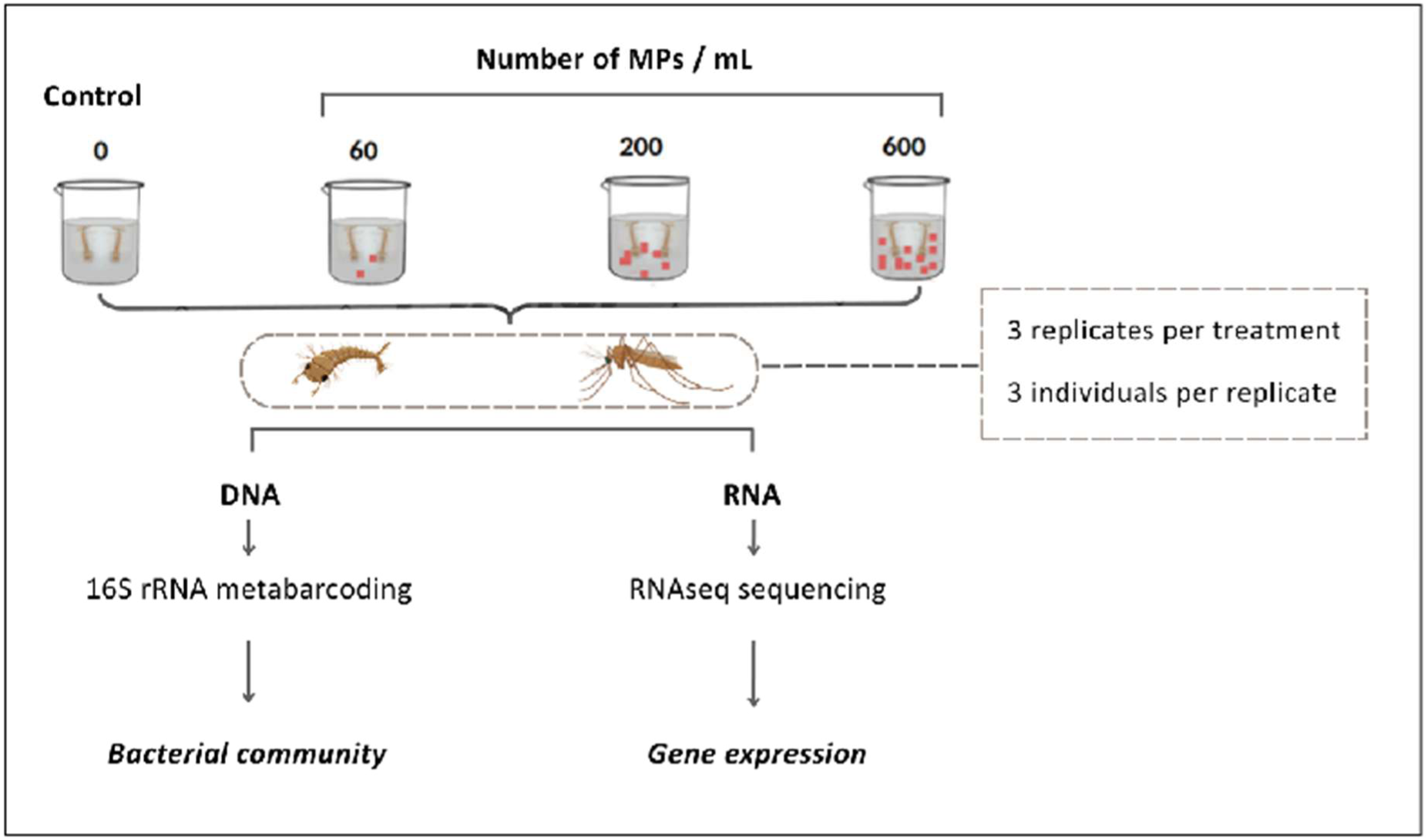
– Overview of the microplastic (MP) exposure experiment and downstream molecular analyses. Mosquito larvae were exposed to increasing MP concentrations (0, 60, 200, and 600 MPs mL⁻¹). For each treatment, three biological replicates were collected, each consisting of pools of three individuals from the same beaker. Fourth-instar larvae (L4) and newly emerged adult females were sampled for simultaneous DNA and RNA extraction. Bacterial community composition was characterized using 16S rRNA metabarcoding, while host gene expression was analyzed by RNA-seq. In addition, remaining adult females from each treatment were dissected to collect midguts for 16S rRNA metabarcoding analysis.

### DNA and RNA extraction and sequencing

Total DNA and RNA from whole larvae and females were simultaneously extracted using the Quick DNA/RNA Miniprep Plus kit (Zymo Research) following the manufacturer’s instructions with slight modifications. Samples were lysed with proteinase K digestion for 2 hours at room temperature. DNA and RNA were separated using the dual-column system and eluted with DNase/RNase-free water using a volume of 75 µL for RNA and 150 µL for DNA. For RNA extraction, an on-column DNase I treatment (15 min at room temperature) was performed to remove genomic DNA contamination. DNA from adult female midguts and dissection negative controls were extracted using the DNeasy Blood & Tissue kit (Qiagen), following the manufacturer’s instructions with an elution volume of 50 µL. Each extraction series incorporated two negative controls: one with kit components alone and a second in which a sterile piston was pre-soaked in the lysis buffer.

For RNA-seq analyses, library preparation and sequencing were performed by the GeT-PlaGe platform (Génopole Toulouse). Polyadenylated transcripts were enriched by poly-A selection. Next-generation sequencing (NGS) was performed in paired-end mode on an AVITI sequencer (Element Biosciences) **(Liu et al., 2024)**.

For microbiota profiling, DNA extracted from larvae, adult whole-body and adult midgut samples was amplified using hypervariable V3-V4 region of the 16S rRNA gene by PCR using universal bacterial primers **(Flores et al., 2023; Juma et al., 2020)**: 341F (5′-CCTACGGGNGGCWGCAG-3′) and 805R (5′-GACTACHVGGGTATCTAATCC-3′). Primers were designed with Illumina adapter sequences (GenSeq) and heterogeneity spacers (N, NN, or NNN) to increase sequence diversity during sequencing. PCR reactions were performed in a final volume of 25 µL containing 1 µL of DNA template (< 30 ng/µL), 5 µL of Phusion HF Buffer (5×), 0.5 µL of dNTP mix (10 mM each), 1.25 µL of each primer (10 µM), 0.25 µL of Phusion High-Fidelity DNA Polymerase (2 U/µL, Thermo Fisher Scientific), and nuclease-free water to volume. Thermal cycling conditions consisted of an initial denaturation at 98°C for 30 s, followed by 30 cycles of denaturation at 98°C for 10 s, annealing at 57°C for 30 s, and extension at 72°C for 1 min 15 s, with a final extension at 72°C for 10 min. All negative controls (dissection controls for the midguts, DNA extraction control and a PCR negative control), along with a positive control (ZymoBIOMICS Microbial Community DNA Standard, from Zymo Research; **Figure S2**), underwent identical amplification procedures. Library preparation and paired-end sequencing (2 × 300 bp) were performed on an Illumina MiSeq system at the GenSeq platform (Montpellier, France).

### Transcriptomic data analysis

RNA-seq reads in FASTQ format were obtained from the GeT-PlaGe platform. Quality control was performed using FastQC v0.12.1 **(Andrews, 2010)**, and reports were aggregated with MultiQC v1.9 **(Ewels et al., 2016)**. Adapter sequences and low-quality bases were removed using fastp v0.20.1 **(Chen et al., 2018)** with default parameters (qualified_quality_phred Q15). Trimmed reads were aligned to the *Cx. quinquefasciatus* reference genome (JHB2020, version 58, VectorBase; NCBI assembly accession: GCF_015732765.1; **Ryazansky et al., 2024**) using HISAT2 v2.2.1 **(Kim et al., 2019)**. Resulting SAM files were converted to BAM format, sorted, and indexed using SAMtools v1.19.2 **(Danecek et al., 2021)**. The genome annotation file (GFF3 format) was converted to GTF format using AGAT v1.5.0 **(Dainat, 2020)**. Gene-level read counts were generated using featureCounts from the Subread package v2.0.6 **(Liao et al., 2014)**. Differential expression analysis was performed separately for larvae and adult females using DESeq2 v1.46.0 **(Love et al., 2014)** with default parameters in R v4.4.1. Genes were considered differentially expressed with an absolute log₂ fold change (Log2FC) > 1 and an adjusted p-value (FDR) < 0.05, using Benjamini-Hochberg correction. For functional enrichment analysis, VectorBase gene identifiers were converted to UniProt IDs as described in **Wei et al. (2024)**, to address incomplete genome annotation. Gene Ontology (GO) enrichment analyses for Biological Process (BP), Molecular Function (MF) and Cellular Component (CC) categories were conducted using ShinyGO v0.77 **(Ge et al., 2020)** with Fisher’s exact test and an FDR cutoff of 0.05.

### Microbiota diversity and composition analyses

Sequencing reads from the 16S rRNA gene obtained from whole individual mosquitoes, dissected midguts and controls were first quality-checked using FastQC v0.12.1 **(Andrews et al., 2021)** and primer-trimmed with Cutadapt v5.0 **(Gong et al., 2011)**. Reads were then processed using the FROGS pipeline v4.1.0 on the Galaxy Platform **(Escudié et al., 2018)**. In our study, paired-end reads were demultiplexed and merged using FLASH v1.2.11 **(Magoč & Salzberg, 2011)** with a maximum mismatch rate of 0.1, and sequences were filtered to retain lengths between 380 and 460 bp. Amplicon sequence variants (hereafter, ASVs) were inferred using Swarm v3.0.0 **(Mahé et al., 2014, 2021)** with an aggregation distance of one nucleotide. Chimeric sequences were detected and removed de novo using VSEARCH v2.17.0 **(Rognes et al., 2016)**, as the singletons (ASVs represented by only one count in the dataset). Taxonomic assignment of ASVs was performed using the naive Bayesian Ribosomal Database Project Classifier algorithm **(Wang et al., 2007)** with the SILVA 139.1 database as reference. ASVs for which the total number of sequences in negative control samples represented more than 50% of the total sequences observed in true samples were discarded, and further contaminant filtering was conducted using the decontam R package **(Davis et al., 2018)**. ASVs assigned to mitochondria and chloroplasts were removed from the dataset. At last, a standardized threshold of 10 counts was applied across all sample types to remove false-positive ASVs, corresponding to 0.005% of the total abundance in the dataset with the lowest sequencing depth (i.e., larvae samples).

Alpha diversity metrics (Shannon, Simpson, Chao1, Faith’s Phylogenetic Diversity (hereafter, PD), and core abundance) were estimated using the vegan **(Oksanen et al., 2025),** phyloseq **(McMurdie & Holmes, 2013)**, picante **(Kembel et al., 2010)** and microbiome **(Lahti & Shetty, 2017)** R packages, respectively. Beta diversity metrics (Bray–Curtis and weighted UniFrac distances) were computed using the microbiome R package from a phyloseq object transformed to relative abundance, in which ASVs were aggregated at the genus level into operational taxonomic units (hereafter, OTU). Based on previous studies **(Juma et al., 2020)**, we hypothesize that bacterial communities differ depending on the stage of the mosquito’s development, but also depending on the sampling scale (whole organism vs. dissected organ). Consequently, prior to investigating the impact of MP exposure, significant differences in alpha diversity and beta diversity among sample types (i.e. larvae, whole females, midgut females) were tested by Kruskal-Wallis’ tests (stats R package: **Lahti and Shetty, 2017**; **R Core Team, 2023**), supported by Dunn tests (FSA R package: **Ogle et al., 2025**; **R Core Team, 2023**), and permutational multivariate analysis of variance (hereafter, PERMANOVA, 9,999 permutations) coupled with permutational analysis of multivariate dispersions (hereafter, BETADISPER) (vegan R package), respectively. Beta diversity was assessed using Principal Coordinates Analysis (PCoA). To avoid a confounding effect with MP exposure, only unexposed individuals were considered. Based on these results (see below), each dataset was subsequently analyzed separately.

Microbial community analyses were conducted to evaluate the impact of MP exposure across treatment groups (exposed vs. unexposed) and concentrations (0, 60, 200, 600 MPs/mL). Alpha diversity values were compared between unexposed and exposed samples using the Wilcoxon-Mann-Whitney test, across concentration categories using the Kruskal-Wallis test (stats R package), and in relation to continuous concentration using Spearman’s rank correlation (stats R package). Beta diversity differences were evaluated by PERMANOVA, with homogeneity of dispersion verified by BETADISPER and PCoA ordination. Linear discriminant analysis was used to compute effect sizes (LDA scores) using the microbiomeMarker R package **(Segata et al., 2011)** to identify genus-level OTUs significantly associated with exposure groups. In parallel, differential abundance was assessed using log2 fold changes calculated with the DESeq2 R package **(Love et al., 2014)**. Comparisons were performed for multiple group contrasts, including unexposed vs. exposed samples, unexposed vs. individual concentration categories, and pairwise comparisons between concentration categories. The ASVs shared across treatments were identified using the ComplexUpset R package. Targeted analyses were performed for Wolbachia and Elizabethkingia OTUs to further characterize exposure-associated patterns. Differences in prevalence between unexposed and exposed groups were assessed using Fisher’s exact test. Among infected samples, relative abundances were compared between exposure groups using the Wilcoxon-Mann-Whitney test and correlated with MP concentration using Spearman’s rank correlation. Log2 fold changes were additionally calculated. For OTUs highlighted as significant by LDA analysis, descriptive assessments were conducted, including visualization of prevalence, mean relative abundance ± standard error (hereafter, SE), and association between relative abundance and MP concentration. Log2 fold changes were likewise estimated.

## Results

### RNA sequencing data description

The proportion of bases with a Phred quality score ≥30 (Q30) ranged from 89.4% to 91.9% in larvae and from 92.4% to 94.7% in adult females. Raw reads ranged from 28.80 to 64.61 million per larval sample and from 37.32 to 74.30 million per adult female sample. Alignment rates to the *Cx. quinquefasciatus* reference genome ranged from 70.0 to 87.9% for larvae and from 87.9 to 96.0% for adult females. GC content varied from 37.5 to 47.6% in larvae and from 24.8 to 30.7% in adult females (reference genome: 35.21%), reflecting differential expression of genes with variable GC content according to developmental stage. The summarized sequencing data and alignment information are reported in **Table S1**.

### Gene expression in larvae

Differential expression analysis (DESeq2, False Discovery Rate **-** FDR < 0.05, |log₂FC| > 1) identified a limited number of differentially expressed genes (DEGs) in larvae exposed to MPs, with a non-monotonic dose-dependent response: 5 DEGs at 60 MPs/mL, 8 DEGs at 200 MPs/mL, and 7 DEGs at 600 MPs/mL (**Figure 2A**). At the lowest concentration (60 MPs/mL), four genes were downregulated, including antimicrobial peptides (*Cecropin-A, Defensin-C*), a lysosomal enzyme (*Lysozyme c-1*), and an endopeptidase inhibitor (CD109), while a non-coding RNA (CQUJHB006533) was highly upregulated (log₂FC = 6.09, FDR <0.05). At 200 MPs/mL, three genes were upregulated and five were downregulated, including the General Odorant Binding Protein (GOBP 56d-like) and genes involved in transmembrane transport (Na-K-Cl cotransporter), cytoskeleton organization (Actin-87E), and chitin degradation (Endochitinase). The long non-coding RNA (lncRNA), CQUJHB006533, also remained upregulated (log₂FC = 5.58; **Figure 2B**). At the highest concentration (600 MPs/mL), seven genes were differentially expressed, six of which were downregulated. This condition was characterized by marked repression of immune system genes, including the same antimicrobial peptides observed at 60 MPs/mL as well as an immune receptor (*Ficolin-1*). The lncRNA CQUJHB006533 showed the strongest upregulation at this concentration (log₂FC = 6.89, FDR <0.05) (**Figure 2C**).

**Figure 2.**
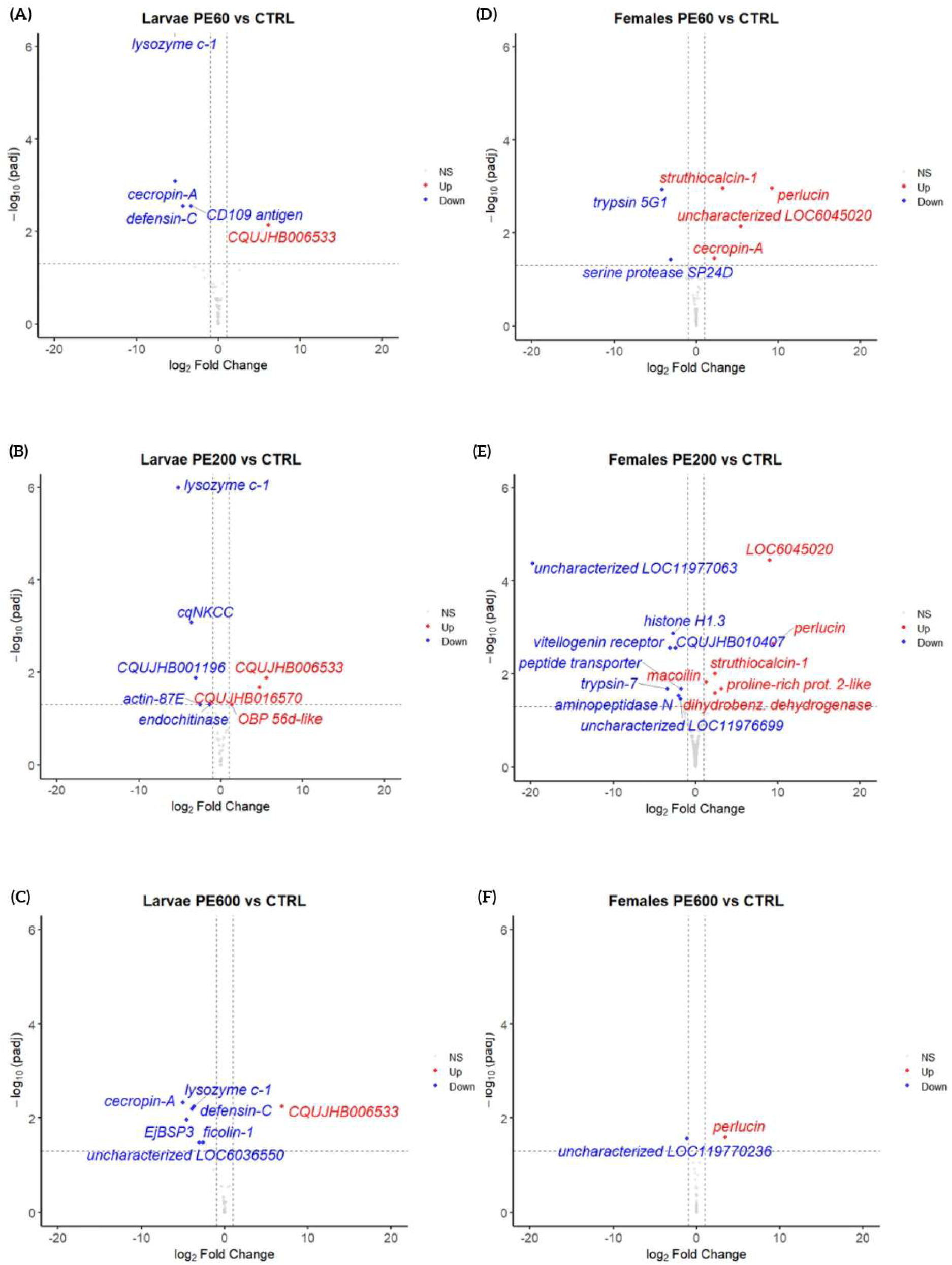
– Volcano plots of differential gene expression in *Cx. quinquefasciatus* larvae and adult females exposed to polyethylene MPs. Differential expression analyses comparing each microplastic concentration (60, 200 and 600 MPs mL⁻¹) to the control condition are shown for larvae (A-C) and newly emerged females (D-F). The x-axis represents log₂ fold change and the y-axis shows −log₁₀ adjusted p-values. Red and blue points indicate significantly up– and down-regulated genes, respectively.

### Gene expression in adult females

In adult females, MP exposure also induced a non-monotonic dose-dependent response: 6 DEGs at 60 MPs/mL, 14 DEGs at 200 MPs/mL and only 2 DEGs at 600 MPs/mL. At 60 MPs/mL, four genes were upregulated: an antimicrobial peptide (*Cecropin-A*), calcium-binding proteins (*Struthiocalcin-1, Perlucin*), and an uncharacterized protein (CQUJHB016620) showing homology to a putative secreted protein from salivary glands. Two serine proteases (*Trypsin 5G1, Serine protease SP24D*) were downregulated (**Figure 2D**). The intermediate concentration (200 MPs/mL) generated the strongest transcriptomic response with 14 DEGs. The six upregulated genes were mainly involved in biomineralization and calcium binding (Perlucin, Struthiocalcin-1), as well as enzymatic activities, whereas the eight downregulated genes were associated with neuronal signal transduction, transmembrane transport, chromatin regulation, and endocytosis. The gene CQUJHB016620, already identified at 60 MPs/mL, remained upregulated. Several uncharacterized genes showed homologies to conserved proteins of the *Culex pipiens* complex, suggesting potential functions in Endoplasmic Reticulum (ER)-associated degradation pathways (ERAD) and ER-Golgi trafficking (**Figure 2E**). The highest concentration (600 MPs/mL) induced only 2 DEGs: Perlucin (upregulated, as at the two other concentrations) and an uncharacterized protein (CQUJHB010486, downregulated) (**Figure 2F**). All DEGs and their log2FC values are reported in **Table S2**.

### Unexposed larvae and females exhibit different bacterial community patterns

A total of 62 samples were examined, including 12 larvae, 12 whole females and 38 female midguts. To ensure the reliability of our results, we included 11 negative controls to monitor for potential contamination during the experiment. Sequencing performance was assessed using a positive control, whose composition included all expected bacterial genera (representing 99.59% of the total abundance), although their relative abundances varied among some genera (**Figure S2**). We obtained 5,902,127 raw sequences from these samples, of which 3,330,994 passed quality filtering (56.44%). While the total sequence counts differed among sample types (**Table S3**), the average number of quality-filtered sequences per sample was comparable and sufficient to support downstream analyses (mean ± SE; larvae: 45,387 ± 6,600 reads; whole females: 58,220 ± 5,874; female midguts: 54,759 ± 3,198). Rarefaction analyses indicated that sequencing depth adequately captured taxonomic richness across sample categories (**Figure S3A**). Larvae exhibited a higher mean number of ASVs (mean ± SE: 240 ± 40 ASVs) compared to females (whole females: 121 ± 25; female midguts: 146 ± 18) (**Table S3**). Further analyses conducted exclusively on unexposed samples (i.e. controls at 0 MPs/mL) revealed significant differences among sample types in estimated species richness (Chao1; Kruskal–Wallis’ test, p-value = 0.034), as well as on the dominance structure and phylogenetic diversity of the bacterial communities (Faith’s PD, p-value = 0.010; core abundance, p-value = 0.032). Larvae showed a significantly higher median Chao1 value compared to females (median + interquartile range, hereafter IQR; larvae: 446.05 + 59.91; whole females: 152.71 + 39.47; female midguts: 132 + 43.92), indicating greater estimated richness. Yet, the taxa constituting larval communities were more phylogenetically clustered and evenly distributed (PD: 7.52 + 0.31; core abundance: 0.001) than those observed in females (whole females, PD: 9.74 + 0.34, core abundance: 0.96 + 0.16; female midguts, PD: 17.41 + 14.14, core abundance: 0.97 + 0.56) (**Figure S3B**). In unexposed whole females, *Wolbachia* (family Ehrlichiaceae) was by far the most abundant genus-level OTU, accounting for an average relative abundance of 85.92 ± 10.55% (**Figure 3A**). A similar pattern was observed in female midguts, where *Wolbachia* represented 71.15 ± 12.29% of the community on average (**Figure S4C**). In contrast, the larval microbiota was not dominated by any specific genus. In this group, *Wolbachia* constituted only 0.09 ± 0.03% of the relative abundance. Instead, at least seven genus-level OTUs exceeded 2% relative abundance per individual, belonging to the families Nevskiaceae, Flavobacteriaceae, Flectobacillaceae, Pseudomonadaceae, Weeksellaceae, Methylophilaceae and Comamonadaceae (**Figure 3A**). The patterns observed for some alpha diversity metrics were consistent with beta diversity indices, which also highlighted significant differences in community structure among sample types (PERMANOVA; WUnifrac, p-value = 0.002; Bray-Curtis, p-value = 0.002) (**Figure S3C**). As a result, we have analyzed each dataset (i.e. whole larvae, whole females and female midguts) separately in subsequent steps. To minimize potential biases related to sampling scale in females (whole organism vs. dissected organ), we focused primarily on comparing the impact of MP exposure between whole-body samples (larvae and whole females).

**Figure 3.**
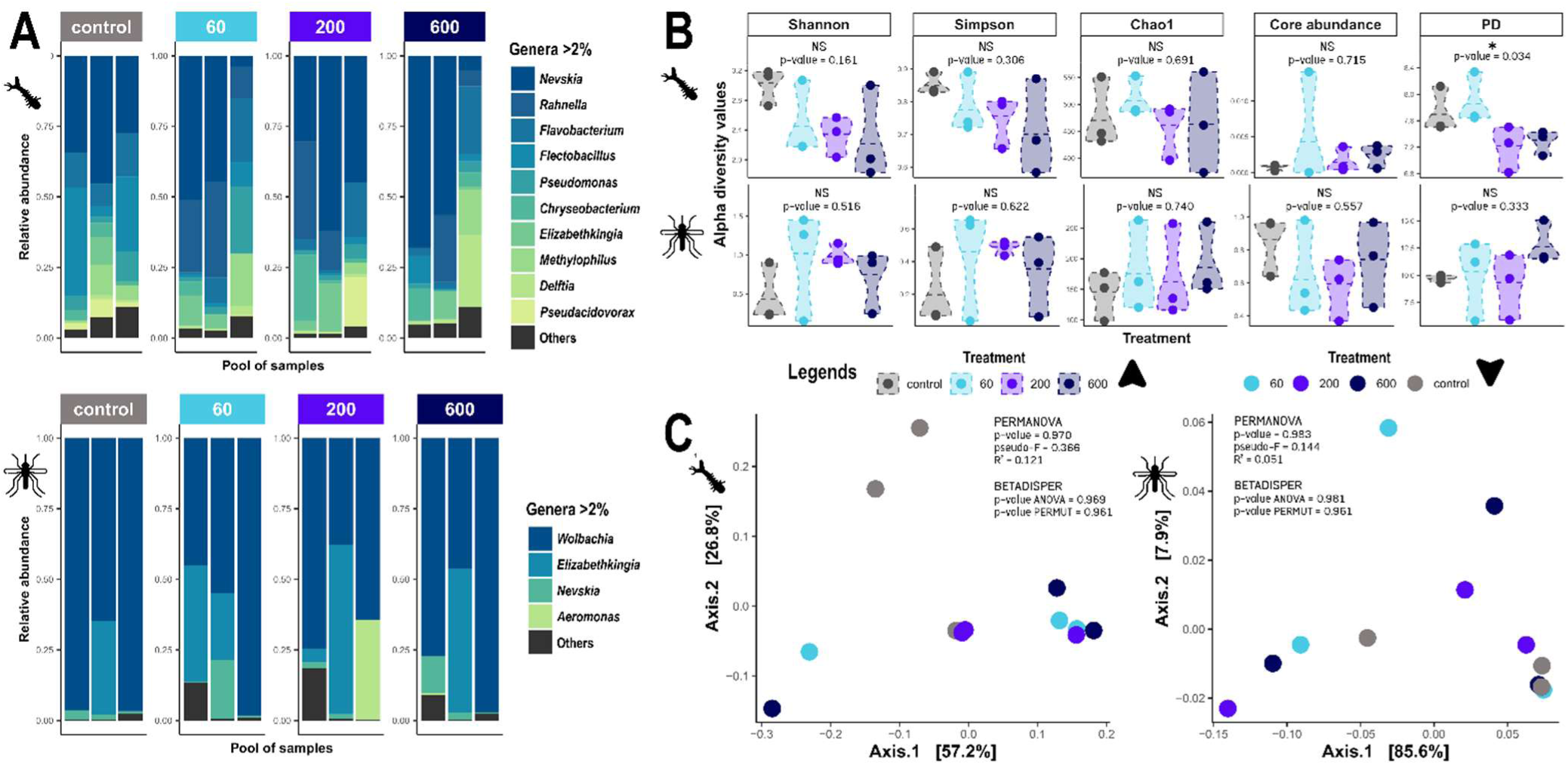
– **A.** Composition plot detailing genus-level OTUs representing at least 2% of abundance data of larvae and whole females, respectively, listed in descending order of abundance. Abundance data are presented as relative abundances and for each MP concentration category. **B.** Alpha diversity metrics and **C.** beta diversity Weighted Unifrac index visualised by PCoA ordination, according to the MP concentration categories for the larvae and whole females, respectively.

### Effects of MP exposure on the microbiota of larvae and females

To assess the effects of MP exposure, we performed comprehensive microbial community analyses across exposure groups and concentration levels. In larvae, we detected a significant effect of exposure status (exposed vs. unexposed) on the Shannon alpha diversity index (Wilcoxon-Mann-Whitney test; p-value = 0.036, effect size = 0.614), indicating that exposure may reduce diversity and potentially compromise overall community stability (median ± IQR: unexposed: 3.129 ± 0.229; exposed: 2.181 ± 0.532) (**Figure S5A**). This finding is further supported by a significant negative correlation between Shannon diversity values and MP concentrations treated as a continuous variable (Spearman’s rank correlation; p-value = 0.023, ρ = −0.648). Similarly, significant differences were observed for the PD index in relation to MP concentrations, both when considered as categorical levels (Kruskal-Wallis’ test; p-value = 0.034, effect size = 0.705) and as a dose gradient (Spearman’s rank correlation; p-value = 0.009, ρ = −0.713). These results suggest shifts in community assembly, with progressively lower phylogenetic dispersion as MP exposure increases (**Figure 3B**; **Figure S5A**). No other significant effects were detected for the remaining alpha diversity indices and variables, despite a visible trend toward decreased Simpson diversity and increased core abundance with increasing MP concentrations.

Likewise, beta diversity metrics did not reveal any significant association with MP exposure status or concentration categories (**Figure 3C**; **Figure S6A**). Interestingly, 39% of ASVs were common to all treatments, 15% were shared among MP treatments and 10% were unique to each treatment (**Figure S7A**). Overall, our results provide some evidence that MP exposure influences the composition and structure of larval bacterial communities. To further characterize this potential effect, we examined whether specific taxa were driving the differences observed among concentration categories (**Figure 4**; **Figure S8C**; **Table 1**). Linear discriminant analysis revealed several genus-level OTUs that were significantly associated with specific exposure groups and displayed marked effect sizes (LDA score > 2), underscoring their discriminatory potential across exposure conditions (**Figure 4A**). Most of the discriminant taxa were enriched in the unexposed group. Notably, *Pseudacidovorax*, *Limnohabitans* and *Sphingorhabdus* showed significantly higher relative abundances in the unexposed group compared with the 600 MPs/mL treatment (Wald test with Benjamini-Hochberg correction, p-value < 0.05; **Figure 4B**), highlighting their contribution to the separation between these conditions. Taxa associated with the unexposed group frequently belonged to the family Comamonadaceae, including *Pseudacidovorax*, an unclassified Comamonadaceae genus, *Alicycliphilus*, *Caenimonas* and *Limnohabitans*. In contrast, *Rahnella*, *Serratia*, *Kluyvera*, *Erwinia* and *Pantoea* were significantly enriched in the treatment 60 MPs/mL relative to the unexposed group (LDA > 2; **Figure 4A**), indicating a concentration-specific response. Differential abundance analysis confirmed these patterns, as these genera were also significantly more prevalent in the 60 MPs/mL group (Wald test with Benjamini-Hochberg correction, p-value < 0.05; **Figure 4B**). Notably, *Rahnella* and *Serratia* consistently exhibited elevated relative abundances in exposed samples compared with unexposed ones (**Figure S9A**). By contrast, *Erwinia, Kluyvera*, and *Pantoea* were not detected in unexposed samples (**Figure S9A**). Several of these concentration-dependent trends were likewise evident in the broader comparison between exposed and unexposed groups, characterized by increased abundances of *Rahnella*, *Serratia*, *Kluyvera*, *Erwinia*, and *Pantoea*, alongside a decrease in *Alicycliphilus* under MP exposure (**Figure S9C**). No significant changes in either relative abundance or prevalence were observed for *Wolbachia* or *Elizabethkingia* OTUs across MP exposure conditions (**Figure S9A**; **Table S4**).

**Figure 4.**
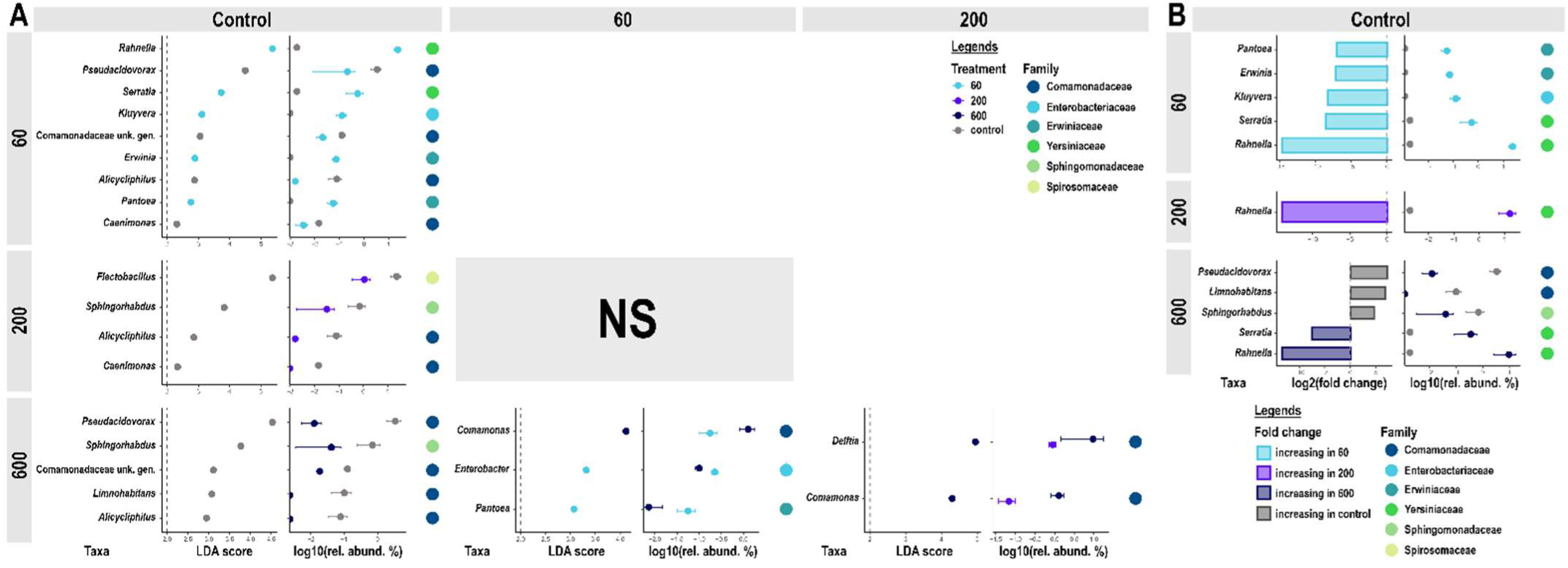
– **A.** Linear discriminant analyses performed for multiple group contrasts, including unexposed vs. individual concentration categories and pairwise comparisons between concentration categories, for the larvae. LDA scores are detailed alongside mean relative abundance per OTU and condition (as log10 values). For each OTU, we precise its taxonomic family. **B.** Differential abundance of genus-level OTUs between the unexposed group and each MP concentration category, for the larvae. We display only OTU with a p-value < 0.05 for the Wald test (including a Benjamini-Hochberg correction). Differential abundance, presented as log2(fold change), are detailed alongside mean relative abundance per OTU and condition (as log10 values). For each OTU, we precise its taxonomic family.

**Table 1.**
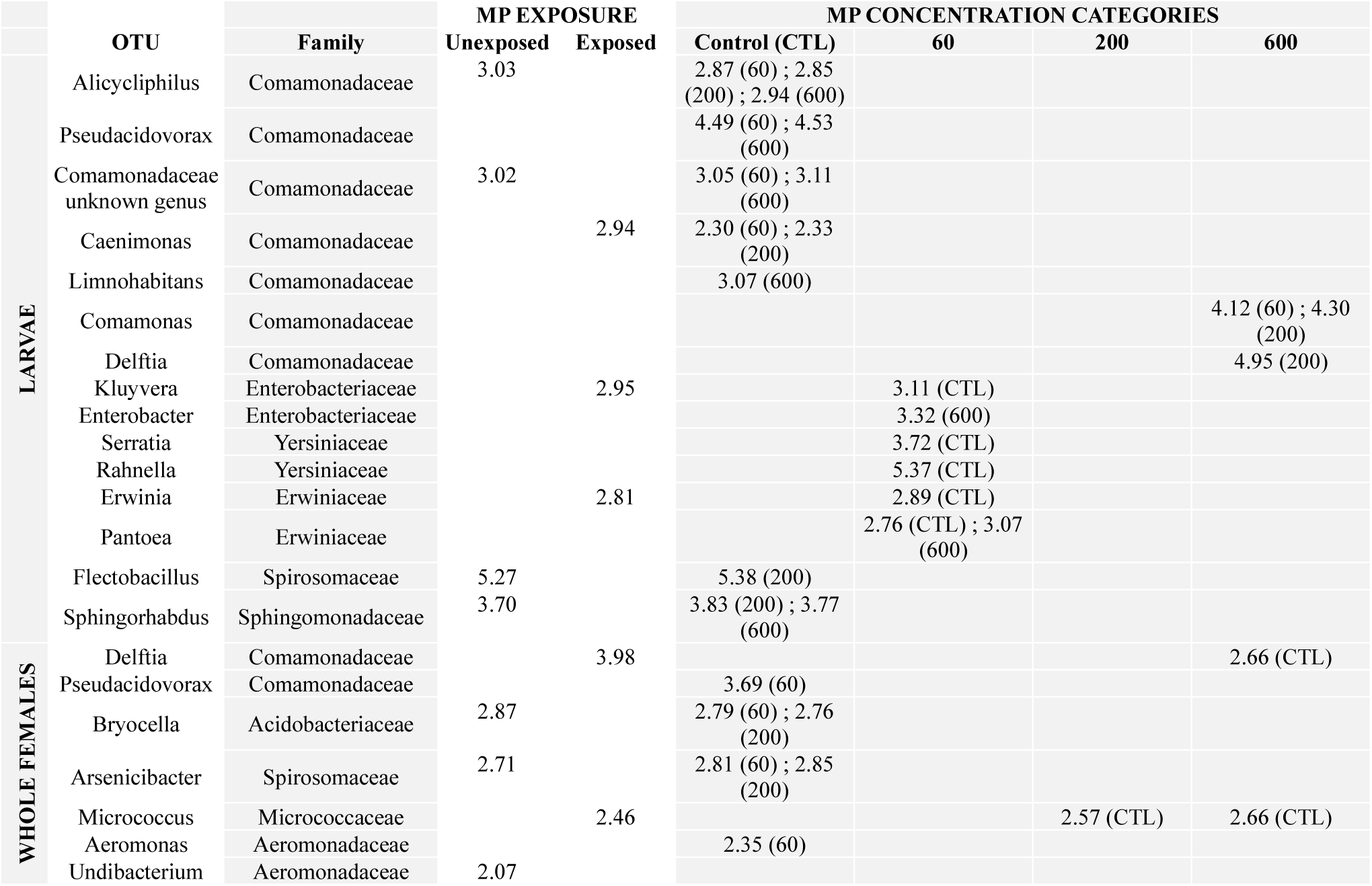
– LDA scores for each discriminant OTU determined during comparisons between levels of exposure to MP and between MP concentration categories. For each OTU, we precise its taxonomic family.

In whole females, MP exposure did not exhibit a clear impact on bacterial communities, nor in terms of species richness or community structure (**Figure 3**; **Figure S5B**; **Figure S6B**). No significant changes were detected in any alpha diversity or beta diversity indices. This lack of significant impact is also observed in female midguts (**Figure S4**). In contrast to results in larvae, 16% of ASVs were common to all treatments, 5% were shared among MP treatments and 50.5% were unique to each treatment (**Figure S7B**). Complementary linear discriminant analyses were performed to identify specific genus-level OTUs associated with exposed groups. Most of the highlighted OTUs were more abundant in the unexposed group, a trend that appears to be primarily driven by their absence from the exposed conditions (**Figure S8A**; **Table 1**). Interestingly, the *Micrococcus* OTU emerged as discriminant for the 200 MPs/mL and 600 MPs/mL treatments compared with controls. Although it did not reach significance in the 60 MPs/mL-specific comparison, a closer inspection indicates that *Micrococcus* OTU *w*as absent in unexposed samples yet consistently present across all exposure treatments (**Figure S9B**). A similar pattern is apparent for the *Delftia* and *Chryseobacterium* OTUs (**Figure S9B**). In addition, the *Chryseobacterium* OTU exhibited significantly higher abundance in the group 60 MPs/mL (**Figure S8B**). Finally, both *Delftia* and *Micrococcus* OTUs were revealed as discriminant taxa in the overall exposed vs. unexposed analysis (**Figure S8D**), further supporting their association with MP exposure. Similar to larvae, no significant changes in either relative abundance or prevalence were observed for *Wolbachia* or *Elizabethkingia* OTUs across MP exposure conditions in whole females (**Figure S9B**; **Table S4**).

Comparison of the two sample types (larvae vs. females) revealed that two specific OTUs were shared: *Pseudacidovorax* and *Delftia*, both members of the family Comamonadaceae (**Table 1**). In both larvae and whole females, *Pseudacidovorax* exhibited a decreasing trend in relative abundance with increasing MP concentration (larvae: log₂ fold change unexposed vs. exposed = −3.87; whole females: −1.69) (**Figure S9**). However, these associations did not reach statistical significance (Spearman’s rank correlation; larvae: p = 0.206, ρ = −0.438; whole females: p = 0.514, ρ = −0.251). In contrast, the tendencies observed for *Delftia* differed between sample types. In larvae, we noted a non-significant increasing trend in relative abundance (log₂(fold change) unexposed vs. exposed = 1.99; Spearman’s rank correlation: p = 0.185, ρ = 0.41). Conversely, in whole females, *Delftia* was absent from the unexposed group and showed a non-significant decreasing trend across exposure levels (Spearman’s rank correlation: p = 0.092, ρ = −0.68) (**Figure S9**).

## Discussion

This study aimed to explore the effects of exposure to MPs at three different concentrations on gene expression and the midgut microbiota in larvae and adult females of *Cx. quinquefasciatus*. We showed that, despite the small number of replicates, some genes were differentially expressed in a stage-specific manner following exposure. In addition, bacterial communities in exposed larvae showed reduced species diversity as MP concentration increased whereas no clear impact on bacterial communities was detected in whole females, either in terms of species richness or community structure. Although linear discriminant analysis identified several bacterial genus-level OTUs significantly associated with specific exposure groups, no clear pattern emerged in terms of community structure in larvae. This study was exploratory by design and its results should therefore be interpreted accordingly. Nevertheless, the observed trends are consistent and warrant further investigation.

### Microplastics affect immune-related gene expression and microbiota in larvae

Direct MP exposure in larvae induced a non-monotonic dose-response pattern on differentially expressed genes (DEGs). At 60 and 600 MPs/mL, DEGs were predominantly associated with immune function, including the downregulation of antimicrobial peptides (Cecropin-A and Defensin-C) and the innate defence enzyme Lysozyme c-1 relative to unexposed larvae. At the intermediate concentration of 200 MPs/mL, DEGs were instead mainly associated with ion transport (Na-K-Cl cotransporter) and cuticle-related processes. Several mechanisms for non-monotonic dose-response effects have been proposed, first in relation to hormones and endocrine disrupting chemicals **(Vandenberg et al., 2012)**, and more recently, after exposure to microplastics **(Sun et al., 2021)**. In the case of MPs, this non-monotonic dose response may reflect concentration-dependent differences in particle retention and bioavailability, where smaller or less aggregated particles may be retained longer in tissues and therefore exerting distinct biological effects **(Jeong et al., 2016)**.

Beyond these concentration-dependent differences, the downregulation of immune-related genes at 60 and 600 MPs/mL warrants specific attention. **Khudhair and Abbood (2026)** reported that exposure of *Ae. aegypti* and *Ae. albopictus* larvae to 1µm polystyrene MPs at concentrations of 1000 and 10 000 MP/µL did not induce significant up or downregulation of Toll or IMD pathway genes, including Defensin A and Cecropin A, despite causing significant gut microbiota disruption. This suggests that MP exposure in mosquito larvae does not always trigger canonical immune activation. MPs could instead induce an indirect effect on immune-related gene expression, for example, in response to a general metabolic stress. Indeed, **Malafaia et al. (2020)** showed that short exposure of *Cx. quinquefasciatus* larvae to the same PE MPs as in our study, at low concentration (around 5 MPs/mL), induced changes indicative of energy metabolism disruption and oxidative stress, suggesting that MPs exposure can constrain metabolic resources in this species. Under conditions of metabolic stress, immune gene expression may be secondarily reduced, given that immune activation is known to be energetically costly in insects **(Schmid-Hempel, 2005).** By contrast, **Doria et al. (2025)** reported immune activation alongside downregulation of metabolic genes in larvae of the freshwater insect *Chironomus riparius* following chronic MPs exposure, highlighting that the direction of immune gene modulation may vary across species, exposure conditions and particle characteristics. Whether the downregulation we observe reflects direct interference of MPs with immune signalling pathways, a consequence of metabolic constraints, or a combination of both, remains to be determined.

The differential expression of osmoregulation and cuticle-related genes at 200 MPs/mL may reflect a response to physical or epithelial stress induced by MPs ingestion at this concentration, given the critical importance of hydromineral balance for aquatic larvae **(Bradley, 1987)**. A comparable pattern has been reported in damselfly larvae (*Ischnura elegans*), where MP exposure upregulated metabolic and cellular stress pathways while immune gene modulation remained limited **(Sun et al., 2025),** suggesting that osmoregulatory and metabolic adjustments may represent a widespread primary response to MPs in aquatic insect larvae.

Regarding MP effects on larval microbiota, MPs appeared to alter bacterial community structure and composition by reducing species richness and phylogenetic dispersion. In addition, *Rahnella* and *Serratia* ASVs were more abundant in MP-exposed larvae, although this increase reached statistical significance only in the 60 MPs/mL treatment. *Serratia* was also identified as a discriminant taxon following exposure to bioplastic MPs in *Daphnia magna* **(Carrillo et al., 2025)**. Interestingly, both *Rahnella* and *Serratia* may be involved in postembryonic development in arthropods. **Valzania et al. (2018)** showed that gut microbiota regulates larval growth in mosquitoes by inducing a hypoxia signal, which, in turn, activates hypoxia-induced transcription factors and other processes required for growth. Similarly, **Liu et al. (2024)** showed that the gut microbiota promoted host development in the red turpentine beetle (*Dendroctonus valens*) by regulation of D-glucose transport from the gut to the fat body. In particular, *Rahnella aquatilis*, *Serratia liquefaciens,* and *Pseudomonas* sp.7, the dominant taxa in the red turpentine beetle, were able to rescue the block on D-glucose transport in their set-up, possibly through the induction of hypoxia and secretion of riboflavin in the gut **(Liu et al., 2024)**. Together, these findings raise the possibility that MPs exposure may indirectly affect larval metabolism and development by altering the relative abundance of bacterial taxa such as *Rahnella* and *Serratia*. *Serratia* spp. may also be involved in plastic degradation, as they are enriched in the gut microbiota of the greater wax moth, *Galleria mellonella* (Lepidoptera), a ‘plastivore’ species capable of degrading polyethylene and polystyrene **(Ruiz Barrionuevo et al., 2022)**.

### Microplastics alter immune-related gene expression but induce no effect on microbiota in adult females

In adult females collected 24-48h after emergence, and therefore exposed to MPs through the larval stage only, an upregulation of immune-related genes was observed, including Cecropin-A, with the most pronounced transcriptomic response at 200 MPs/mL (14 DEGs). On the other hand, the long non-coding RNA (lncRNA) CQUJHB006533 that was consistently upregulated across all MP concentrations in larvae (log₂FC ≈ 6.1-6.9), was not differentially expressed in newly emerged adult females. LncRNAs are known to participate in epigenetic regulation, notably through chromatin remodeling and transcription factor sequestration **(Belavilas-Trovas et al., 2023; Moure et al., 2022)**. Such mechanisms could produce persistent changes in gene expression states that outlast the initial transcriptional signals. We therefore hypothesize that MP-induced lncRNA activity in larvae contributed to epigenetic modifications, which may in turn have shaped transcriptional programs in adult females and may potentially take apart in the immune gene upregulation observed here.

In contrast to what was observed in larvae, MP exposure had no detectable effect on microbiota structure or composition in adult females. In a similar experimental setup, **Edwards et al. (2023)** reported an increase in *Elizabethkingia* abundance in *Ae. aegypti* and *Ae. albopictus* females following larval exposure to PS MPs, pattern that was not observed in our study. This discrepancy may reflect interspecific differences, as *Elizabethkingia* was more abundant in those species than in *Cx. quinquefasciatus*.

### MPs exposure may trigger carry-over effects

Females were not directly exposed to MPs as adults, which raise the question of how larval MPs exposure could be associated with differential gene expression after metamorphosis. Two main hypotheses can be proposed. First, residual MPs in adult tissues could directly influence gene expression, as suggested by evidence that MPs acquired during the larval stage can be retained in the adult mosquitoes **(Al-Jaibachi et al., 2018)**. Second, larval exposure may have exerted carry-over effects on adults. According to the definition given by **Moore and Martin (2019)**, a carry-over effect is a phenotype, which is not directly observed at the stage of exposure, but at a succeeding life stage. Carry-over effects of larval conditions on adult immune phenotype have been documented in insects in other contexts. **Fellous and Lazzaro (2011)** showed in *Drosophila melanogaster* that larval and adult immune gene expression shared common genetic determinants, implying that perturbations affecting larval immunity can have consequences for adult immune phenotypes, suggesting that such cross-stage effects can occur in insects.

Although no carry-over effects were observed on the female microbiota, MP-induced shifts in larval bacterial communities may nevertheless influence other biological processes associated with the microbiota, including immune function, metabolism or life history traits. To date, most studies investigating MPs exposure in mosquitoes have focused on adult traits following larval exposure **(Al-Jaibachi et al., 2019; Edwards et al., 2023; Griffin et al., 2023; Malafaia et al., 2020; Misser et al., 2025; Thormeyer & Tseng, 2023)**. For instance, **Ezeakacha and Yee (2019)** observed a carry-over effect of larval rearing temperatures on the fecundity of the mosquito *Ae. albopictus* (*i.e.* decreased fecundity with increased larval rearing temperature). Given the limited retention of MPs after metamorphosis, most of the observed effects in adults are likely to represent carry-over effects rather than direct consequences of MPs persistence **(Al-Jaibachi et al., 2019)**. Moreover, **Aryaprema et al. (2025)** showed that exposure to 1 μm and 30 μm PS MPs (10000 and 100,000 MPs/mL) affected larval development, adult body size, blood-feeding behaviour and fecundity in a species-specific way in *Cx. quinquefasciatus* and *Anopheles quadrimaculatus*.

While direct mosquito mortality following MP exposure has been reported **(Griffin et al., 2023)** with potential consequences for population dynamics, our findings suggest that MP-induced microbiota dysbiosis and alterations in immune gene regulation may also influence mosquito populations.

### Limitations and future directions

Although our findings provide novel insights, they should be interpreted in light of several methodological limitations. The small number of biological replicates (n = 3), combined with the high inter-individual variability characteristic of mosquito transcriptomes, likely explains, at least in part, the low number of DEGs detected under each exposure condition. Similarly, our sampling size may have limited our ability to detect subtle but yet genuine shifts in microbiota community structure. Increasing the number of replicates in future experiments would improve statistical power and facilitate the detection of additional biologically relevant patterns. The use of pristine MPs also limits the ecological realism of our study, as environmental MPs are typically colonized by biofilms and can adsorb co-contaminants such as heavy metals and pesticides **(Luo et al., 2022)**, which may further complicate interactions with the larval microbiota. Complementary experiments using environmentally aged MPs collected from urban parks and stormwater drain systems are currently underway which will greatly improve ecological relevance. Future studies should also determine whether residual MPs persist in adult tissues using appropriate physicochemical analyses, thereby allowing direct residual exposure to be distinguished from carry-over effects. Finally, our microbiota analyses were restricted to bacteria. Incorporating fungal ITS sequencing and viral metagenomics would provide a more comprehensive understanding of host-microbiome interactions under MPs exposure.

As mosquitoes are worldwide disease vectors and are increasingly exposed to MPs present throughout aquatic environments, future studies should focus on determining how the MP-induced effects may affect the vector competence of mosquitoes, and thus their ability to transmit diseases. The partial immune gene modulation observed in adult females in our study, despite their lack of direct exposure, raises the possibility that larval MP exposure could influence adult susceptibility to pathogen infection **(Loiseau & Sorci, 2022)**. Studies in *Ae. albopictus* have shown that adult MP exposure can reduce Zika virus transmission through a combination of physical adsorption of viral particles and immune pathway modulation **(Li et al., 2025)**, demonstrating that such effects are biologically plausible. Future studies should therefore examine transcriptomic and immune responses in adult females following infectious blood meals at key time points throughout the extrinsic incubation period and integrate these analyses with pathogen quantification and transmission efficiency assays.

## Acknowledgements

We thank the Vectopole platform team for providing the laboratory space to perform the experiments. We also thank the GenSeq platform staff for their support. The authors acknowledge the ISO 9001 certified IRD itrop HPC (member of the South Green Platform) at IRD Montpellier for providing HPC resources that have contributed to the research results reported within this paper (https://bioinfo.ird.fr/; http://www.southgreen.fr). We thank Florence Nono Almeida for our team discussions. This work is part of the Zone Atelier Camargue.

## Data, scripts, code, and supplementary information availability

The transcriptomic and metabarcoding raw reads are deposited under The BioProject PRJNA1522685 on NCBI (https://www.ncbi.nlm.nih.gov/).

All the codes used for this study are stored in a GitHub repository (https://github.com/mariebuysse/Microplastics_gene_microbiota), joined to a Zenodo DOI (XXX) [pending].

Supplementary material is available online on bioRxiv along with the main manuscript.

## References

1. Al-Jaibachi, R., Cuthbert, R. N., & Callaghan, A. (2018). Up and away: Ontogenic transference as a pathway for aerial dispersal of microplastics. Biology Letters, 14(9), 20180479. 10.1098/rsbl.2018.0479

2. Al-Jaibachi, R., Cuthbert, R. N., & Callaghan, A. (2019). Examining effects of ontogenic microplastic transference on Culex mosquito mortality and adult weight. Science of The Total Environment, 651, 871–876. 10.1016/j.scitotenv.2018.09.236

3. Andrews, S. (2010). FastQC: A quality control tool for high throughput sequence data [Logiciel]. http://www.bioinformatics.babraham.ac.uk/projects/fastqc/

4. Andrews, T. S., Kiselev, V. Y., McCarthy, D., & Hemberg, M. (2021). Tutorial: Guidelines for the computational analysis of single-cell RNA sequencing data. Nature Protocols, 16(1), 1–9. 10.1038/s41596-020-00409-w

5. Apte-Deshpande, A. D., Paingankar, M. S., Gokhale, M. D., & Deobagkar, D. N. (2014). Serratia odorifera mediated enhancement in susceptibility of Aedes aegypti for chikungunya virus. 139. https://pmc.ncbi.nlm.nih.gov/articles/PMC4140042/

6. Arif, Y., Mir, A. R., Zielinski, P., Hayat, S., & Bajguz, A. (2024, avril). Microplastics and nanoplastics: Source, behavior, remediation, and multi-level environmental impact. Journal of Environmental Management, 356. 10.1016/j.jenvman.2024.120618

7. Aryaprema, V. S., Silva, J. D., Qualls, W. A., & Xue, R.-D. (2025). Effects of polystyrene microplastic ingestion on development, adult fitness, and reproductive succes of Culex quinquefasciatus and Anopheles Quadrimaculatus. Journal of the Florida Mosquito Control Association, 72. 10.32473/jfmca.72.1.139355

8. Bahia, A. C., Dong, Y., Blumberg, B. J., Mlambo, G., Tripathi, A., BenMarzouk-Hidalgo, O. J., Chandra, R., & Dimopoulos, G. (2014). Exploring *Anopheles* gut bacteria for *Plasmodium* blocking activity. Environmental Microbiology, 16(9), 2980–2994. 10.1111/1462-2920.12381

9. Banaee, M., Multisanti, C. R., Impellitteri, F., Piccione, G., & Faggio, C. (2025). Environmental toxicology of microplastic particles on fish: A review. Comparative Biochemistry and Physiology Part C: Toxicology & Pharmacology, 287, 110042. 10.1016/j.cbpc.2024.110042

10. Belavilas-Trovas, A., Tastsoglou, S., Dong, S., Kefi, M., Tavadia, M., Mathiopoulos, K. D., & Dimopoulos, G. (2023). Long non-coding RNAs regulate Aedes aegypti vector competence for Zika virus and reproduction. PLOS Pathogens, 19(6), e1011440. 10.1371/journal.ppat.1011440

11. Bradley, T. (1987). Physiology of Osmoregulation in Mosquitoes. Annual Review of Entomology, 32, 439–462. https://www.annualreviews.org/content/journals/10.1146/annurev.en.32.010187.002255

12. Cansado-Utrilla, C., Zhao, S. Y., McCall, P. J., Coon, K. L., & Hughes, G. L. (2021). The microbiome and mosquito vectorial capacity: Rich potential for discovery and translation. Microbiome, 9(1), 111. 10.1186/s40168-021-01073-2

13. Carrasco-Navarro, V., Muñiz-González, A.-B., Sorvari, J., & Martínez-Guitarte, J.-L. (2021). Altered gene expression in Chironomus riparius (insecta) in response to tire rubber and polystyrene microplastics. Environmental Pollution, 285, 117462. 10.1016/j.envpol.2021.117462

14. Carrillo, M. P., Vila-Costa, M., & Barata, C. (2025). Micro-bioplastic impact on gut microbiome, cephalic transcription and cognitive function in the aquatic invertebrate Daphnia magna. Environmental Pollution, 382, 126690. 10.1016/j.envpol.2025.126690

15. Charlton-Howard, H. S., Bond, A. L., Rivers-Auty, J., & Lavers, J. L. (2023). ‘Plasticosis’: Characterising macro– and microplastic-associated fibrosis in seabird tissues. Journal of Hazardous Materials, 450, 131090. 10.1016/j.jhazmat.2023.131090

16. Chen, S., Zhou, Y., Chen, Y., & Gu, J. (2018). fastp: An ultra-fast all-in-one FASTQ preprocessor. Bioinformatics, 34(17), i884-i890. 10.1093/bioinformatics/bty560

17. Cuthbert, R. N., Al-Jaibachi, R., Dalu, T., Dick, J. T. A., & Callaghan, A. (2019). The influence of microplastics on trophic interaction strengths and oviposition preferences of dipterans. Science of The Total Environment, 651, 2420–2423. 10.1016/j.scitotenv.2018.10.108

18. Dainat, J. (2020). AGAT: Another Gff Analysis Toolkit to handle annotations in any GTF/GFF format (Version Version v0.7.0) [Logiciel]. 10.5281/zenodo.3552717

19. Danecek, P., Bonfield, J. K., Liddle, J., Marshall, J., Ohan, V., Pollard, M. O., Whitwham, A., Keane, T., McCarthy, S. A., Davies, R. M., & Li, H. (2021). Twelve years of SAMtools and BCFtools. GigaScience, 10(2), giab008. 10.1093/gigascience/giab008

20. Davis, N. M., Proctor, D. M., Holmes, S. P., Relman, D. A., & Callahan, B. J. (2018). Simple statistical identification and removal of contaminant sequences in marker-gene and metagenomics data. Microbiome, 6(1), 226. 10.1186/s40168-018-0605-2

21. Doria, H. B., Sohal, N., Feldmeyer, B., & Pfenninger, M. (2025). Size over substance: Microplastic particle size drives gene expression and fitness loss in a freshwater insect. Aquatic Toxicology, 284, 107386. 10.1016/j.aquatox.2025.107386

22. Edwards, C.-C., McConnel, G., Ramos, D., Gurrola-Mares, Y., Arole, K. D., Green, M. J., Cañas-Carrell, J. E., & Brelsfoard, C. L. (2023). *Microplastic ingestion perturbs the microbiome of Aedes albopictus and Aedes aegypti* [Preprint]. In Review. 10.21203/rs.3.rs-2535203/v1

23. Escudié, F., Auer, L., Bernard, M., Mariadassou, M., Cauquil, L., Vidal, K., Maman, S., Hernandez-Raquet, G., Combes, S., & Pascal, G. (2018). FROGS: Find, Rapidly, OTUs with Galaxy Solution. Bioinformatics, 34(8), 1287–1294. 10.1093/bioinformatics/btx791

24. Ewels, P., Magnusson, M., Lundin, S., & Käller, M. (2016). MultiQC: Summarize analysis results for multiple tools and samples in a single report. Bioinformatics, 32(19), 3047–3048. 10.1093/bioinformatics/btw354

25. Ezeakacha, N. F., & Yee, D. A. (2019). The role of temperature in affecting carry-over effects and larval competition in the globally invasive mosquito Aedes albopictus. Parasites & Vectors, 12(1), 123. 10.1186/s13071-019-3391-1

26. Fellous, S., & Lazzaro, B. P. (2011). Potential for evolutionary coupling and decoupling of larval and adult immune gene expression. Molecular Ecology, 20(7), 1558–1567. 10.1111/j.1365-294X.2011.05006.x

27. Franzellitti, S., Canesi, L., Auguste, M., Wathsala, R. H. G. R., & Fabbri, E. (2019). Microplastic exposure and effects in aquatic organisms: A physiological perspective. Environmental Toxicology and Pharmacology, 68, 37–51. 10.1016/j.etap.2019.03.009

28. Gabrieli, P., Caccia, S., Varotto-Boccazzi, I., Arnoldi, I., Barbieri, G., Comandatore, F., & Epis, S. (2021). Mosquito Trilogy: Microbiota, Immunity and Pathogens, and Their Implications for the Control of Disease Transmission. Frontiers in Microbiology, 12, 630438. 10.3389/fmicb.2021.630438

29. Gao, H., Cui, C., Wang, L., Jacobs-Lorena, M., & Wang, S. (2020). Mosquito Microbiota and Implications for Disease Control. Trends in Parasitology, 36(2), 98–111. 10.1016/j.pt.2019.12.001

30. Ge, S. X., Jung, D., & Yao, R. (2020). ShinyGO: A graphical gene-set enrichment tool for animals and plants. Bioinformatics, 36(8), 2628–2629. 10.1093/bioinformatics/btz931

31. Gong, T., Hartmann, N., Kohane, I. S., Brinkmann, V., Staedtler, F., Letzkus, M., Bongiovanni, S., & Szustakowski, J. D. (2011). Optimal Deconvolution of Transcriptional Profiling Data Using Quadratic Programming with Application to Complex Clinical Blood Samples. PLoS ONE, 6(11), e27156. 10.1371/journal.pone.0027156

32. Griffin, C. D., Tominiko, C., Medeiros, M. C. I., & Walguarnery, J. W. (2023). Microplastic pollution differentially affects development of disease-vectoring Aedes and Culex mosquitoes. Ecotoxicology and Environmental Safety, 267, 115639. 10.1016/j.ecoenv.2023.115639

33. Hillyer, J. (2010). Mosquito immunity. Adv Exp Med Biol, 708, 218–238. https://www.sciencedirect.com/science/article/abs/pii/S2214574514000376

34. Iannuzzi, Z. (2025). Identification des sources et des voies de transfert de microplastiques dans un hydrosystème urbain. INSA Lyon. https://theses.fr/2025ISAL0123

35. Jeong, C.-B., Won, E.-J., Kang, H.-M., Lee, M.-C., Hwang, D.-S., Hwang, U.-K., Zhou, B., Souissi, S., Lee, S.-J., & Lee, J.-S. (2016). Microplastic Size-Dependent Toxicity, Oxidative Stress Induction, and p-JNK and p-p38 Activation in the Monogonont Rotifer (Brachionus koreanus). Environmental Science & Technology, 50(16), 8849–8857. 10.1021/acs.est.6b01441

36. Juma, E. O., Allan, B. F., Kim, C.-H., Stone, C., Dunlap, C., & Muturi, E. J. (2020). Effect of life stage and pesticide exposure on the gut microbiota of Aedes albopictus and Culex pipiens L. Scientific Reports, 10(1), 9489. 10.1038/s41598-020-66452-5

37. Kembel, S. W., Cowan, P. D., Helmus, M. R., Cornwell, W. K., Morlon, H., Ackerly, D. D., Blomberg, S. P., & Webb, C. O. (2010). Picante: R tools for integrating phylogenies and ecology. Bioinformatics, 26(11), 1463–1464. 10.1093/bioinformatics/btq166

38. Khudhair, I. H., & Abbood, N. M. (2026). Microplastic-Induced Dysbiosis in Aedes aegypti and Aedes albopictus Larvae Without Immune Activation. Journal of Applied Health Sciences and Medicine, 6(3), 7. 10.58614/jahsm632

39. Kim, D., Paggi, J. M., Park, C., Bennett, C., & Salzberg, S. L. (2019). Graph-based genome alignment and genotyping with HISAT2 and HISAT-genotype. Nature Biotechnology, 37(8), 907–915. 10.1038/s41587-019-0201-4

40. Koelmans, A. A., Besseling, E., Wegner, A., & Foekema, E. M. (2013). Plastic as a Carrier of POPs to Aquatic Organisms: A Model Analysis. Environ. Sci. Technol., 47, 7812–7820.

41. Kumar, A., Srivastava, P., Sirisena, P., Dubey, S. K., Kumar, R., Shrinet, J., & Sunil, S. (2018). Mosquito Innate Immunity. Insects, 9(3), 95. 10.3390/insects9030095

42. Lahti, L., & Shetty, S. (2017). Microbiome R package. Bioconductor. 10.18129/B9.bioc.microbiome

43. Lee, Y., Sung, M., Sung, S.-E., Choi, J.-H., Kang, K.-K., Park, J. W., Kim, Y., & Lee, S. (2025). The histopathological and functional consequences of microplastic exposure. Discover Applied Sciences, 7(1), 72. 10.1007/s42452-025-06470-y

44. Li, J., Liu, X., Gao, H., Liang, G., Zhao, T., & Li, C. (2025). Not for nothing, microplastics can (potentially) reduce the risk of mosquito-to-human transmission of arboviruses. Journal of Hazardous Materials, 492, 138166. 10.1016/j.jhazmat.2025.138166

45. Li, J., Liu, X., Liang, G., Gao, H., Guo, S., Zhou, X., Xing, D., Zhao, T., & Li, C. (2024). Microplastics affect mosquito from aquatic to terrestrial lifestyles and are transferred to mammals through mosquito bites. Science of The Total Environment, 917, 170547. 10.1016/j.scitotenv.2024.170547

46. Liao, Y., Smyth, G. K., & Shi, W. (2014). featureCounts: An efficient general purpose program for assigning sequence reads to genomic features. Bioinformatics, 30(7), 923–930. 10.1093/bioinformatics/btt656

47. Limonta, G., Mancia, A., Benkhalqui, A., Bertolucci, C., Abelli, L., Fossi, M. C., & Panti, C. (2019). Microplastics induce transcriptional changes, immune response and behavioral alterations in adult zebrafish. Scientific Reports, 9(1), 15775. 10.1038/s41598-019-52292-5

48. Liu, F., Ye, F., Yang, Y., Kang, Z., Liu, Y., Chen, W., Wang, S., Kou, H., Kang, L., & Sun, J. (2024). Gut bacteria are essential for development of an invasive bark beetle by regulating glucose transport. Proceedings of the National Academy of Sciences, 121(33), e2410889121. 10.1073/pnas.2410889121

49. Liu, S., Obert, C., Yu, Y.-P., Zhao, J., Ren, B.-G., Liu, J.-J., Wiseman, K., Krajacich, B. J., Wang, W., Metcalfe, K., Smith, M., Ben-Yehezkel, T., & Luo, J.-H. (2024). Utility analyses of AVITI sequencing chemistry. BMC Genomics, 25(1), 778. 10.1186/s12864-024-10686-4

50. Loiseau, C., & Sorci, G. (2022). Can microplastics facilitate the emergence of infectious diseases? Science of The Total Environment, 823, 153694. 10.1016/j.scitotenv.2022.153694

51. Love, M. I., Huber, W., & Anders, S. (2014). Moderated estimation of fold change and dispersion for RNA-seq data with DESeq2. Genome Biology, 15(12), 550. 10.1186/s13059-014-0550-8

52. Lu, L., Wan, Z., Luo, T., Fu, Z., & Jin, Y. (2018). Polystyrene microplastics induce gut microbiota dysbiosis and hepatic lipid metabolism disorder in mice. Science of The Total Environment, 631*-632*, 449-458. 10.1016/j.scitotenv.2018.03.051

53. Luo, H., Liu, C., He, D., Xu, J., Sun, J., Li, J., & Pan, X. (2022). Environmental behaviors of microplastics in aquatic systems: A systematic review on degradation, adsorption, toxicity and biofilm under aging conditions. Journal of Hazardous Materials, 423(126915). 10.1016/j.jhazmat.2021.126915

54. Lv, W.-X., Cheng, P., Lei, J.-J., Peng, H., Zang, C.-H., Lou, Z.-W., Liu, H.-M., Guo, X.-X., Wang, H.-Y., Wang, H.-F., Zhang, C.-X., Liu, L.-J., & Gong, M.-Q. (2023). Interactions between the gut micro-community and transcriptome of Culex pipiens pallens under low-temperature stress. Parasites & Vectors, 16(1), 12. 10.1186/s13071-022-05643-7

55. Magoč, T., & Salzberg, S. L. (2011). FLASH: Fast length adjustment of short reads to improve genome assemblies. Bioinformatics, 27(21), 2957–2963. 10.1093/bioinformatics/btr507

56. Mahé, F., Czech, L., Stamatakis, A., Quince, C., De Vargas, C., Dunthorn, M., & Rognes, T. (2021). Swarm v3: Towards tera-scale amplicon clustering. Bioinformatics, 38(1), 267–269. 10.1093/bioinformatics/btab493

57. Mahé, F., Rognes, T., Quince, C., De Vargas, C., & Dunthorn, M. (2014). Swarm: Robust and fast clustering method for amplicon-based studies. PeerJ, 2, e593. 10.7717/peerj.593

58. Malafaia, G., Da Luz, T. M., Guimarães, A. T. B., & Araújo, A. P. C. (2020). Polyethylene microplastics are ingested and induce biochemical changes in Culex quinquefasciatus (Diptera: Culicidae) freshwater insect larvae. ECOTOXICOLOGY AND ENVIRONMENTAL CONTAMINATION, 15, 79–89. 10.5132/eec.2020.01.10

59. McConnel, G., Cuellar, D., Arole, K. D., Dasari, S. S., Green, M. J., Cañas-Carrell, J. E., & Brelsfoard, C. L. (2024). Characterization of microplastics found in mosquito oviposition habitats. Journal of Vector Ecology, 50(1), 39–47. 10.52707/1081-1710-50.1-39

60. McMurdie, P. J., & Holmes, S. (2013). phyloseq: An R Package for Reproducible Interactive Analysis and Graphics of Microbiome Census Data. PLoS ONE, 8(4), e61217. 10.1371/journal.pone.0061217

61. Misser, S., Chen, C., Ismail, A., & Oliver, S. V. (2025). The Effect of Larval Exposure to Plastic Pollution on the Gut Microbiota of the Major Malaria Vector *ANOPHELES ARABIENSIS* Patton (Diptera: Culicidae). Environmental Microbiology Reports, 17(4), e70169. 10.1111/1758-2229.70169

62. Moore, M. P., & Martin, R. A. (2019). On the evolution of carry-over effects. Journal of Animal Ecology, 88(12), 1832–1844. 10.1111/1365-2656.13081

63. Moure, U. A. E., Tan, T., Sha, L., Lu, X., Shao, Z., Yang, G., Wang, Y., & Cui, H. (2022). Advances in the Immune Regulatory Role of Non-Coding RNAs (miRNAs and lncRNAs) in Insect-Pathogen Interactions. Frontiers in Immunology, 13, 856457. 10.3389/fimmu.2022.856457

65. Nguyen, T. T., Park, K., Kim, S., & Kwak, I.-S. (2026). Toxicity of nano– and microplastics on marine and freshwater fish: A multi-scale perspective from organ systems to cells. Water Biology and Security, 5(4), 100534. 10.1016/j.watbs.2025.100534

66. OECD. (2022a). Global Plastics Outlook: Economic Drivers, Environmental Impacts and Policy Options. OECD Publishing. 10.1787/de747aef-en

67. OECD. (2022b). Global Plastics Outlook: Policy Scenarios to 2060. OECD Publishing. 10.1787/aa1edf33-en

68. Ogle, D. H., Doll, J. C., Wheeler, P., & Dinno, A. (2025). FSA: Simple Fisheries Stock Assessment Methods. 10.32614/CRAN.package.FSA

69. Oksanen, J., Simpson, G. L., Blanchet, F. G., Kindt, R., Legendre, P., & Minchin, P. R. (2025). Vegan: Community Ecology Package (Version 2.7-1) [Logiciel].

70. Onyango, M. G., Lange, R., Bialosuknia, S., Payne, A., Mathias, N., Kuo, L., Vigneron, A., Nag, D., Kramer, L. D., & Ciota, A. T. (2021). Zika virus and temperature modulate Elizabethkingia anophelis in Aedes albopictus. Parasites & Vectors, 14(1), 573. 10.1186/s13071-021-05069-7

71. R Core Team. (2023). R: A language and Environùent for Statistical Computing. R Foundation for Statistical Computing. https://www.R-project.org/

72. Rafa, N., Ahmed, B., Zohora, F., Bakya, J., Ahmed, S., Ahmed, S. F., Mofijur, M., Chowdhury, A. A., & Almomani, F. (2024). Microplastics as carriers of toxic pollutants: Source, transport, and toxicological effects. Environmental Pollution, 343, 123190. 10.1016/j.envpol.2023.123190

73. Reid, W. R., Zhang, L., Gong, Y., Li, T., & Liu, N. (2018). Gene expression profiles of the Southern house mosquito *Culex quinquefasciatus* during exposure to permethrin. Insect Science, 25(3), 439–453. 10.1111/1744-7917.12438

74. Rodrigues Dos Santos, C., Pinheiro Drumond, G., Rezende Moreira, V., Valéria De Souza Santos, L., & Cristina Santos Amaral, M. (2023). Microplastics in surface water: Occurrence, ecological implications, quantification methods and remediation technologies. Chemical Engineering Journal, 474, 144936. 10.1016/j.cej.2023.144936

75. Rognes, T., Flouri, T., Nichols, B., Quince, C., & Mahé, F. (2016). VSEARCH: A versatile open source tool for metagenomics. PeerJ, 4, e2584. 10.7717/peerj.2584

76. Ruiz Barrionuevo, J. M., Vilanova-Cuevas, B., Alvarez, A., Martín, E., Malizia, A., Galindo-Cardona, A., De Cristóbal, R. E., Occhionero, M. A., Chalup, A., Monmany-Garzia, A. C., & Godoy-Vitorino, F. (2022). The Bacterial and Fungal Gut Microbiota of the Greater Wax Moth, Galleria mellonella L. Consuming Polyethylene and Polystyrene. Frontiers in Microbiology, 13, 918861. 10.3389/fmicb.2022.918861

77. Ryazansky, S. S., Chen, C., Potters, M., Naumenko, A. N., Lukyanchikova, V., Masri, R. A., Brusentsov, I. I., Karagodin, D. A., Yurchenko, A. A., Dos Anjos, V. L., Haba, Y., Rose, N. H., Hoffman, J., Guo, R., Menna, T., Kelley, M., Ferrill, E., Schultz, K. E., Qi, Y., … Sharakhova, M. V. (2024). The chromosome-scale genome assembly for the West Nile vector Culex quinquefasciatus uncovers patterns of genome evolution in mosquitoes. BMC Biology, 22(1), 16. 10.1186/s12915-024-01825-0

78. Sahai, H., García Valverde, M., Murcia Morales, M., Hernando, M. D., Aguilera Del Real, A. M., & Fernández-Alba, A. R. (2023). Exploring sorption of pesticides and PAHs in microplastics derived from plastic mulch films used in modern agriculture. Chemosphere, 333, 138959. 10.1016/j.chemosphere.2023.138959

79. Schmid-Hempel, P. (2005). Evolutionary Ecology of Insect Immune Defenses. Annual Review of Entomology, 529–551.

80. Segata, N., Izard, J., Waldron, L., Gevers, D., Miropolsky, L., Garrett, W. S., & Huttenhower, C. (2011). Metagenomic biomarker discovery and explanation. Genome Biology, 12(6), R60. 10.1186/gb-2011-12-6-r60

81. Shi, X., Chen, Z., Wei, W., Chen, J., & Ni, B.-J. (2023). Toxicity of micro/nanoplastics in the environment: Roles of plastisphere and eco-corona. Soil & Environmental Health, 1(1), 100002. 10.1016/j.seh.2023.100002

82. Sun, L., Cheng, Z., Wang, M., Wei, C., Liu, H., & Yang, Y. (2025). A multi-levels analysis to evaluate the toxicity of microplastics on aquatic insects: A case study with damselfly larvae (Ischnura elegans). Ecotoxicology and Environmental Safety, 289, 117447. 10.1016/j.ecoenv.2024.117447

83. Sun, T., Zhan, J., Li, F., Ji, C., & Wu, H. (2021). Effect of microplastics on aquatic biota: A hormetic perspective. Environmental Pollution, 285, 117206. 10.1016/j.envpol.2021.117206

84. Thormeyer, M., & Tseng, M. (2023). No Effect of Realistic Microplastic Exposure on Growth and Development of Wild-caught Culex (Diptera: Culicidae) Mosquitoes. Journal of Medical Entomology, 60(3), 604–607. 10.1093/jme/tjad014

85. Valzania, L., Martinson, V. G., Harrison, R. E., Boyd, B. M., Coon, K. L., Brown, M. R., & Strand, M. R. (2018). Both living bacteria and eukaryotes in the mosquito gut promote growth of larvae. PLOS Neglected Tropical Diseases, 12(7), e0006638. 10.1371/journal.pntd.0006638

86. Vandenberg, L. N., Colborn, T., Hayes, T. B., Heindel, J. J., Jacobs, D. R., Jr., Lee, D.-H., Shioda, T., Soto, A. M., vom Saal, F. S., Welshons, W. V., Zoeller, R. T., & Myers, J. P. (2012). Hormones and Endocrine-Disrupting Chemicals: Low-Dose Effects and Nonmonotonic Dose Responses. Endocrine Reviews, 33(3), 378–455. 10.1210/er.2011-1050

87. Wang, K., Li, J., Zhao, L., Mu, X., Wang, C., Wang, M., Xue, X., Qi, S., & Wu, L. (2021). Gut microbiota protects honey bees (Apis mellifera L.) against polystyrene microplastics exposure risks. Journal of Hasardous Materials, 402(123828). 10.1016/j.jhazmat.2020.123828

88. Wang, Q., Garrity, G. M., Tiedje, J. M., & Cole, J. R. (2007). Naïve Bayesian Classifier for Rapid Assignment of rRNA Sequences into the New Bacterial Taxonomy. Applied and Environmental Microbiology, 73(16), 5261–5267. 10.1128/AEM.00062-07

89. Wei, X., Lee, K., Mullassery, N., Dhungana, P., Kang, D. S., & Sim, C. (2024). Transcription profiling reveals tissue-specific metabolic pathways in the fat body and ovary of the diapausing mosquito Culex pipiens. Comparative Biochemistry and Physiology Part D: Genomics and Proteomics, 51, 101260. 10.1016/j.cbd.2024.101260

90. Wu, P., Sun, P., Nie, K., Zhu, Y., Shi, M., Xiao, C., Liu, H., Liu, Q., Zhao, T., Chen, X., Zhou, H., Wang, P., & Cheng, G. (2019). A Gut Commensal Bacterium Promotes Mosquito Permissiveness to Arboviruses. Cell Host & Microbe, 25(1), 101–112.e5. 10.1016/j.chom.2018.11.004

91. Yee, D. A., Kesavaraju, B., & Juliano, S. A. (2004). Larval feeding behavior of three co-occurring species of container mosquitoes. Journal of Vector Ecology: Journal of the Society for Vector Ecology, 29(2), 315. https://pmc.ncbi.nlm.nih.gov/articles/PMC2582444/pdf/nihms76018.pdf

