## Supplementary material for "Microplastic exposure alters immune-related gene expression in *Culex quinquefasciatus* mosquitoes and larval microbiota diversity"

### Supplementary figures

**Supplementary figure 1**

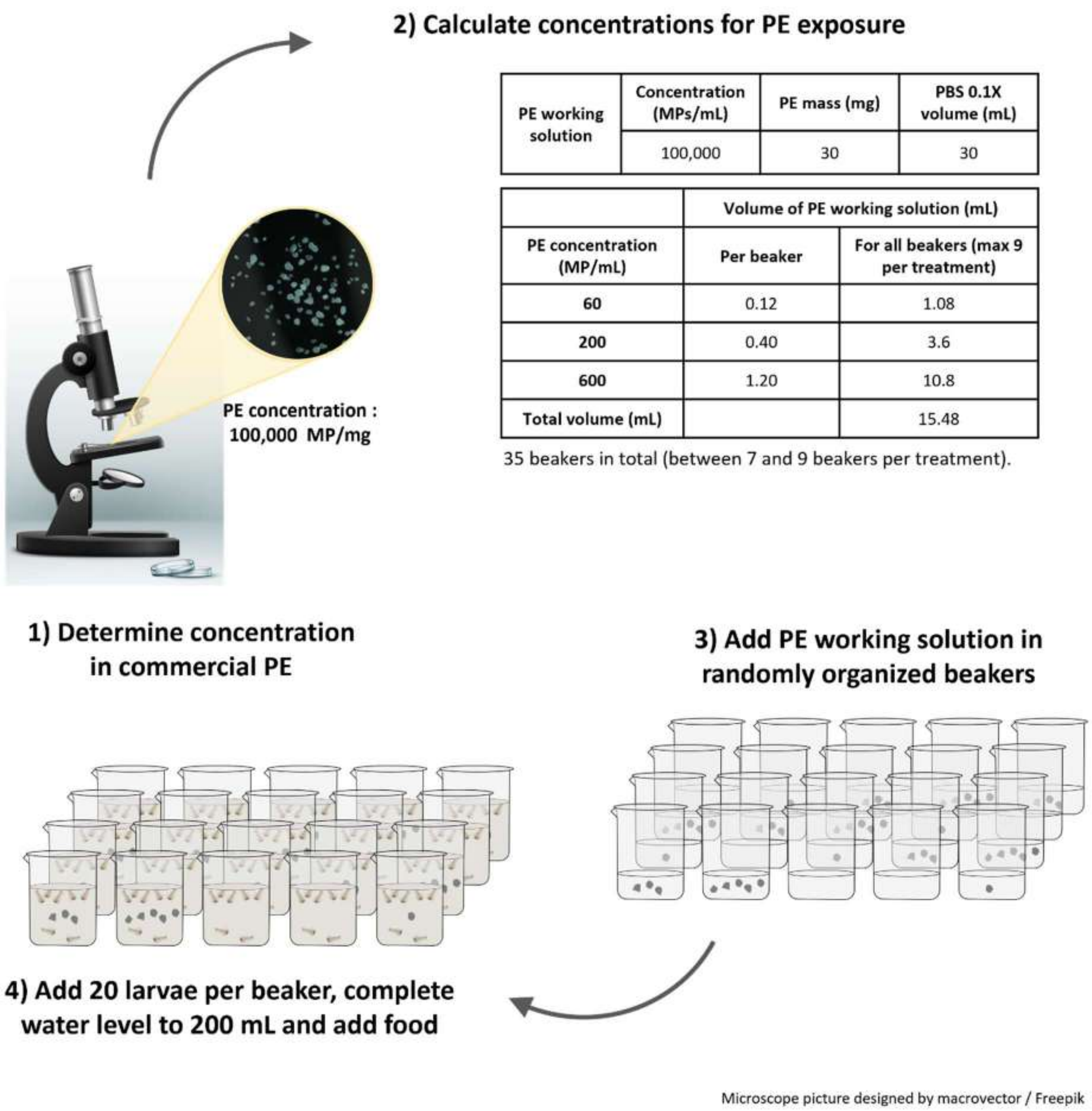

| | Concentration (MPs/mL) | PE mass (mg) | PBS 0.1X volume (mL) |
|---|---|---|---|
| PE working solution | 100,000 | 30 | 30 |

| | Volume of PE working solution (mL) | |
|---|---|---|
| PE concentration (MP/mL) | Per beaker | For all beakers (max 9 per treatment) |
| 60 | 0.12 | 1.08 |
| 200 | 0.40 | 3.6 |
| 600 | 1.20 | 10.8 |
| Total volume (mL) | | 15.48 |

**Figure S1. Description of the experimental protocol used for Culex quinquefasciatus larvae exposure to polyethylene (PE) microplastics (MPs). 1)** First, tests were made to identify the number of PE MPs present in a given mass of the commercial product. A solution with a known concentration of PE MPs (in mg PE MPs/ mL) was prepared, and pictures were made of drops of 1 uL of the solution using a microscope associated with a digital camera. PE MPs could be counted on the pictures and the number of PE MPs per mg of commercial product estimated. **2)** Given an estimated concentration of 100,000 PE MPs/mg, calculations were made to prepare the stock solution for PE exposure. The stock solution was prepared in PBS 0.1X to increase the dispersion of PE MPs in water, while minimising the toxicological effects of using a solvent like ethanol. **3)** The stock solution was distributed in the glass beakers based on the required concentration, and the beakers were randomly numbered to avoid any bias. A small volume of water was put in the beakers prior to adding the stock solution to avoid PE MPs sticking to the glass after evaporation. Control beakers were prepared with 1.20 mL of PBS, which corresponds to the highest volume of stock solution used in the other beakers. **4)** Finally, 20 first instar (L1) or second instar (L2) larvae (less than 48h from egg hatching) were added to each beaker. Reverse osmosis water was added up to 200 mL for each beaker (*N.B.* beakers were filled up to 200 mL regularly during the exposure to avoid an artificial increase of PE concentrations due to water evaporation). And two halves of guinea pig pellets (Safe - Premium Scientific Diet) were added to each beaker.

**Supplementary figure 2**

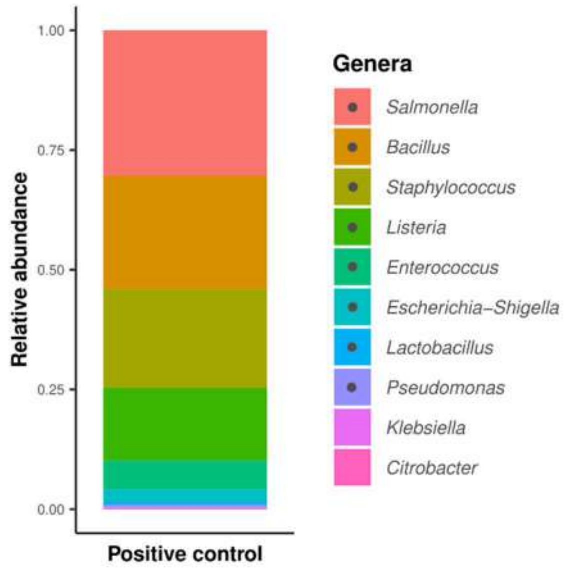

**Figure S2. Composition plot showing all genus-level operational taxonomic units (OTUs) identified in the positive control**, listed in descending order of abundance. Abundance data are presented as relative abundances. Genera expected in the reference material used to generate the positive control are indicated by grey dots.

**Supplementary figure 3**

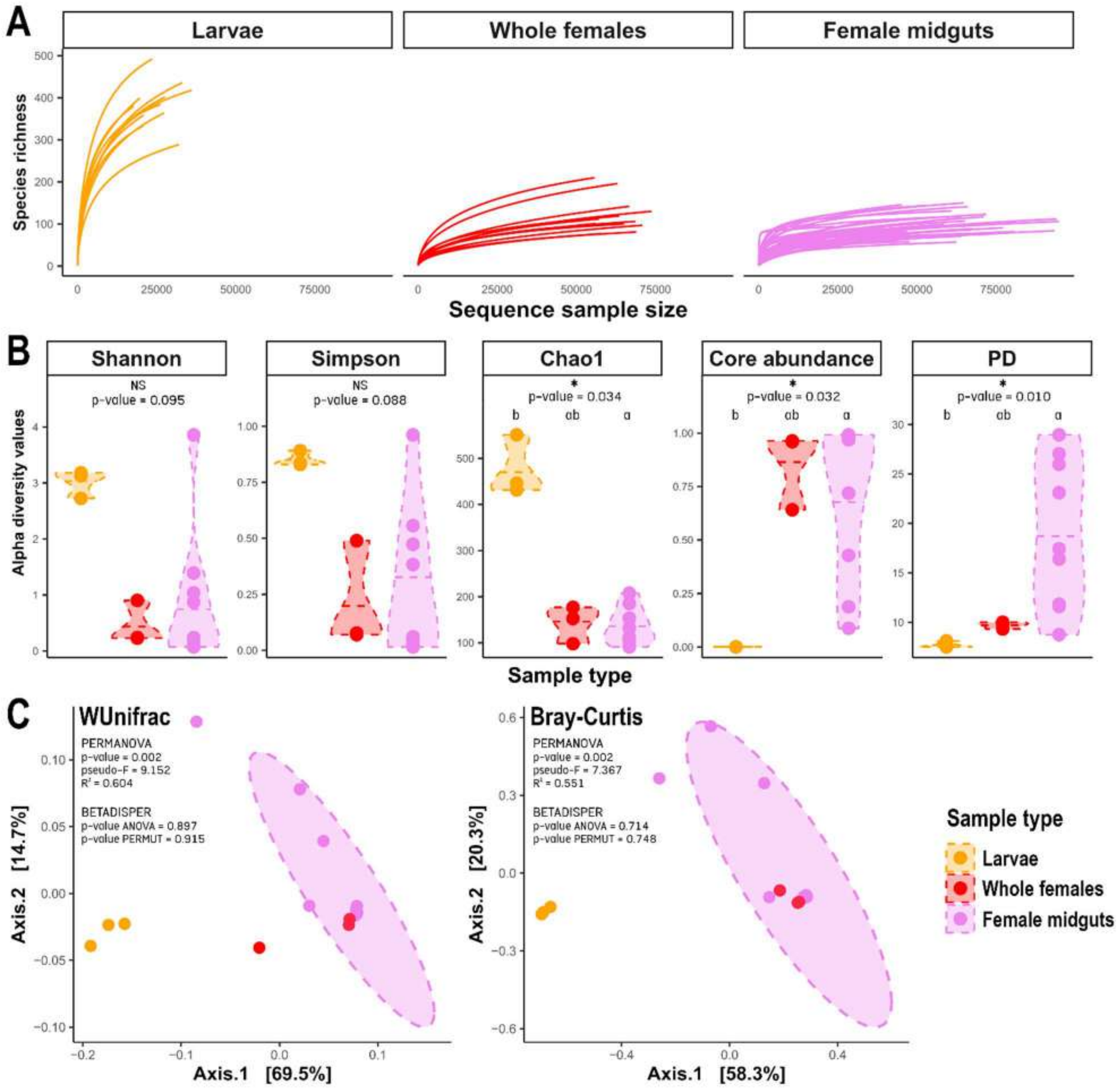

**Figure S3. Rarefaction curves (A), alpha diversity metrics (B), and beta diversity indices (C) visualised by PCoA ordination according to the sample type.**

**Supplementary figure 4**

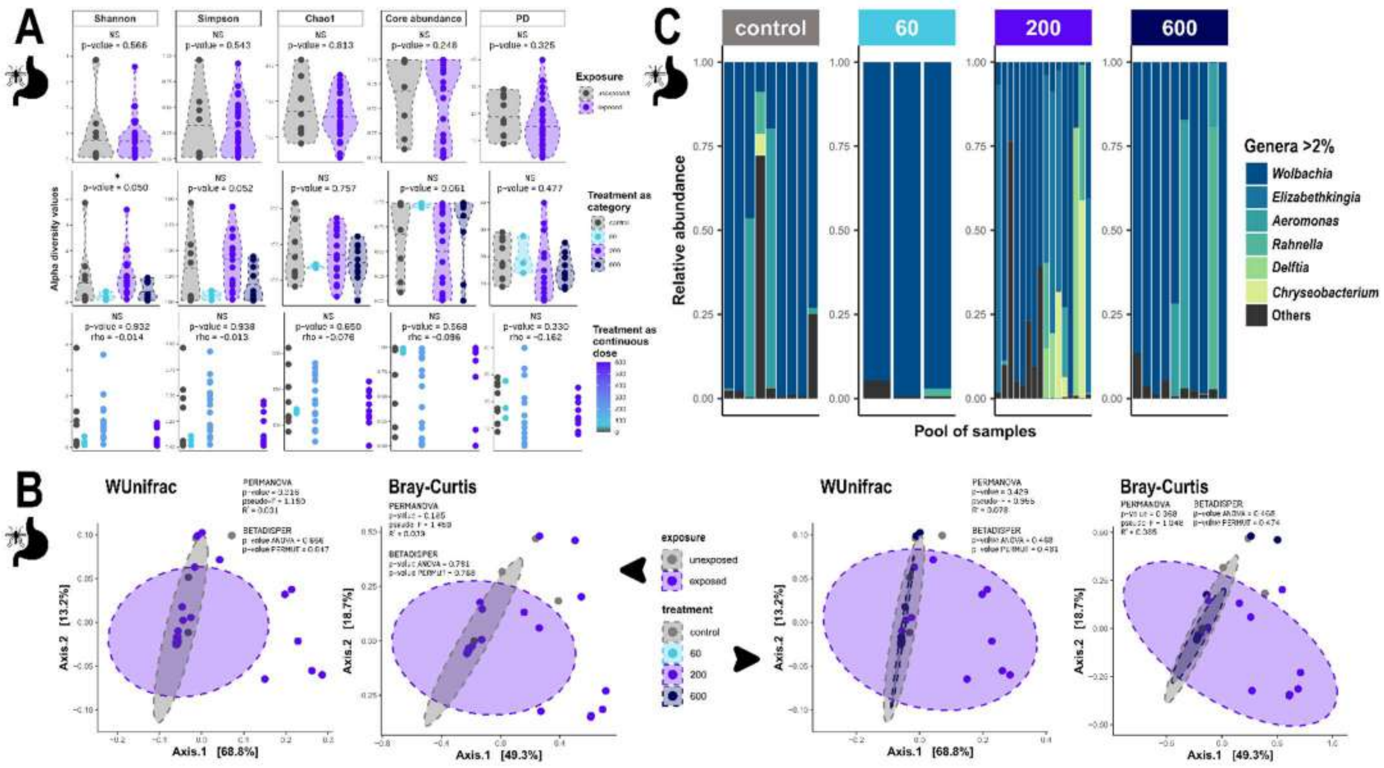

**Figure S4. Diversity metrics and composition plots for metabarcoding of female midguts.** (A) Alpha diversity metrics and (B) beta diversity indices visualised by PCoA ordination, according to the MP exposure status and the MP concentration categories for the female midguts. (C) Composition plot detailing genus-level OTUs representing at least 2% of abundance data of female midguts, listed in descending order of abundance. Abundance data are presented as relative abundances and for each MP concentration category.

*Dissection protocol: To avoid any contamination during dissections, forceps and histological slides were immersed in bleach for 5 min, followed by two rinses in Milli-Q water for 2 min each. Prior to dissection, each mosquito was surface-sterilised by immersion in bleach for 20 seconds, followed by two rinses in Milli-Q water for 1 min each. For each dissection session, negative controls were included by immersing instruments (forceps and a piece of paper towel used to wipe the histological slide) in sterile PBS 1X. Midguts were dissected individually, stored in 1.5 mL Eppendorf tubes containing PBS 1X, and kept at -20°C until DNA extraction.*

**Supplementary figure 5**

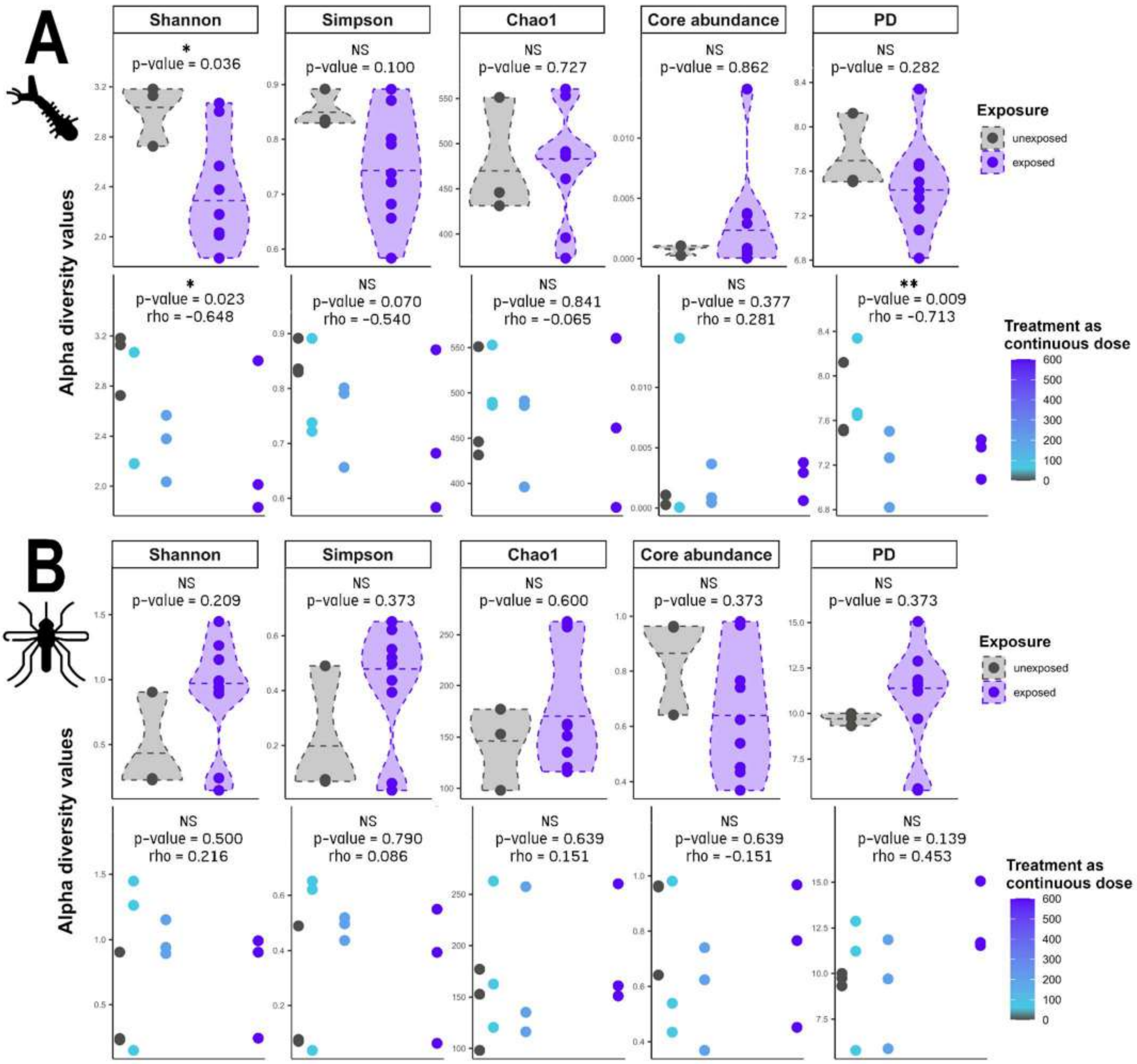

**Figure S5. Alpha diversity metrics according to the MP exposure status and the MP concentration (as continuous doses) for the larvae (A) and the whole females (B).**

**Supplementary figure 6**

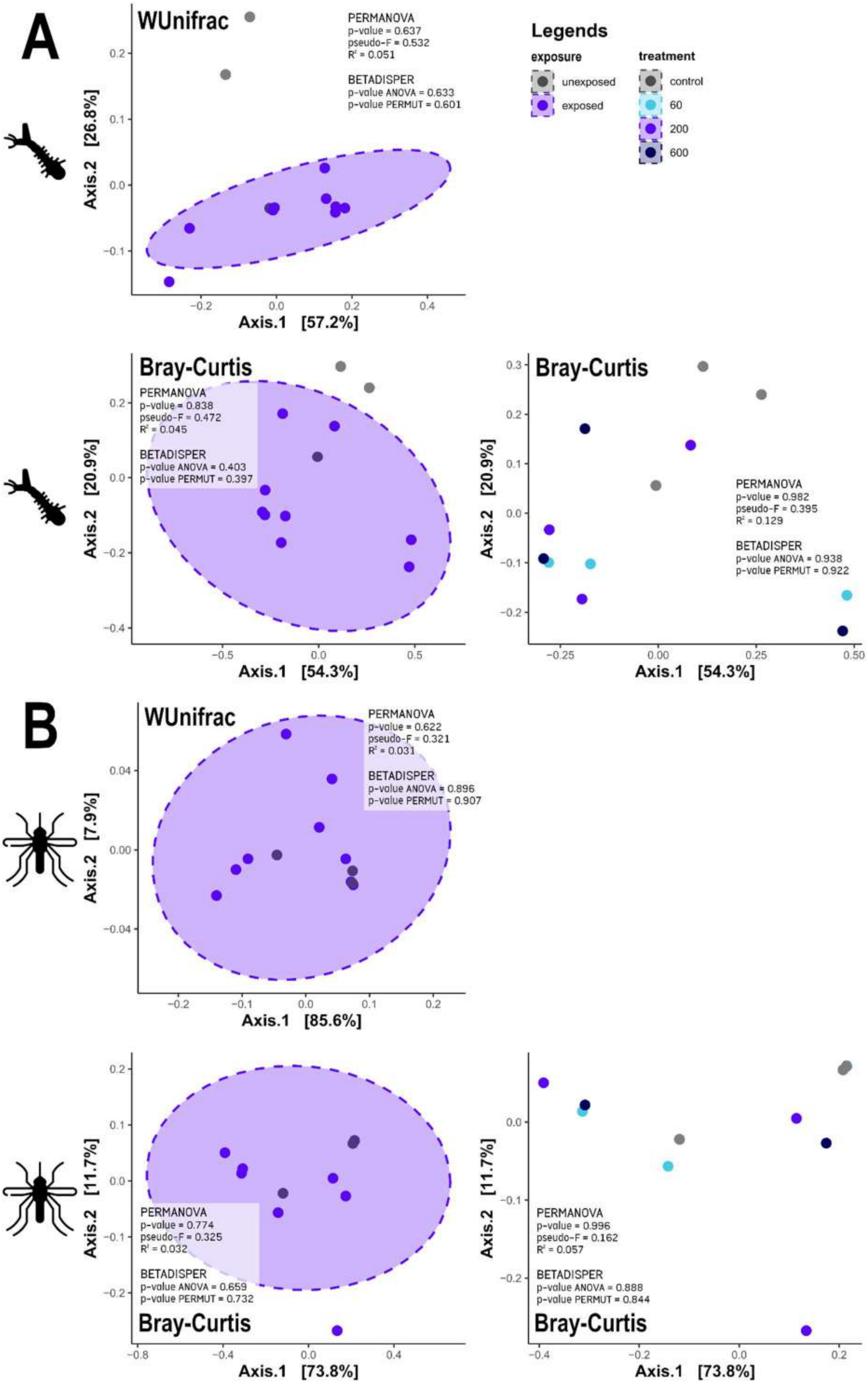

**Figure S6. Beta diversity indices (Weighted Unifrac and Bray-Curtis) visualised by PCoA ordination** according to the MP exposure status and the MP concentration categories for the larvae (A) and the whole females (B).

**Supplementary figure 7**

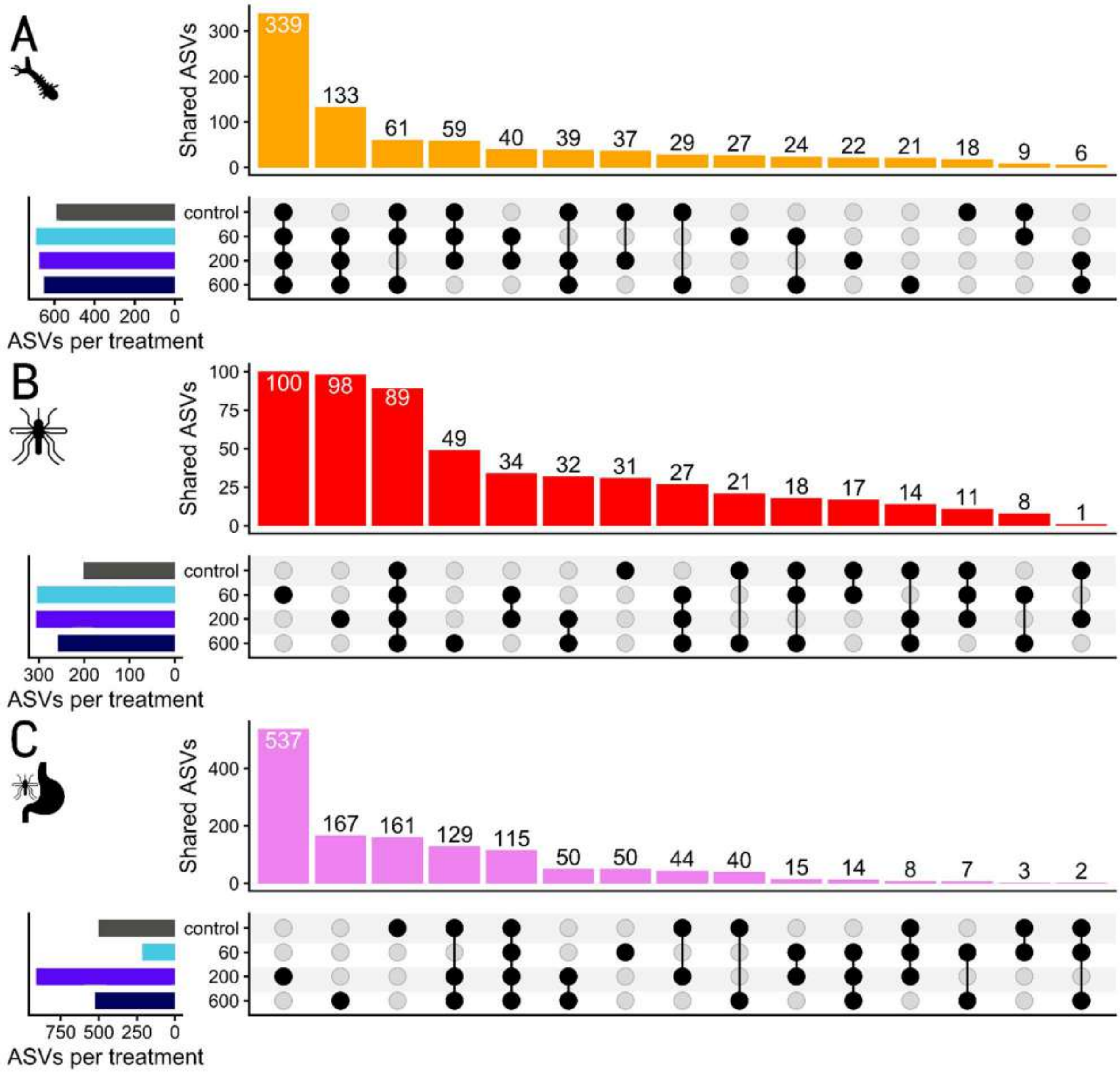

**Figure S7. Shared ASVs across MP treatments** for the larvae (A), the whole females (B), and the female midguts (C).

**Supplementary figure 8**

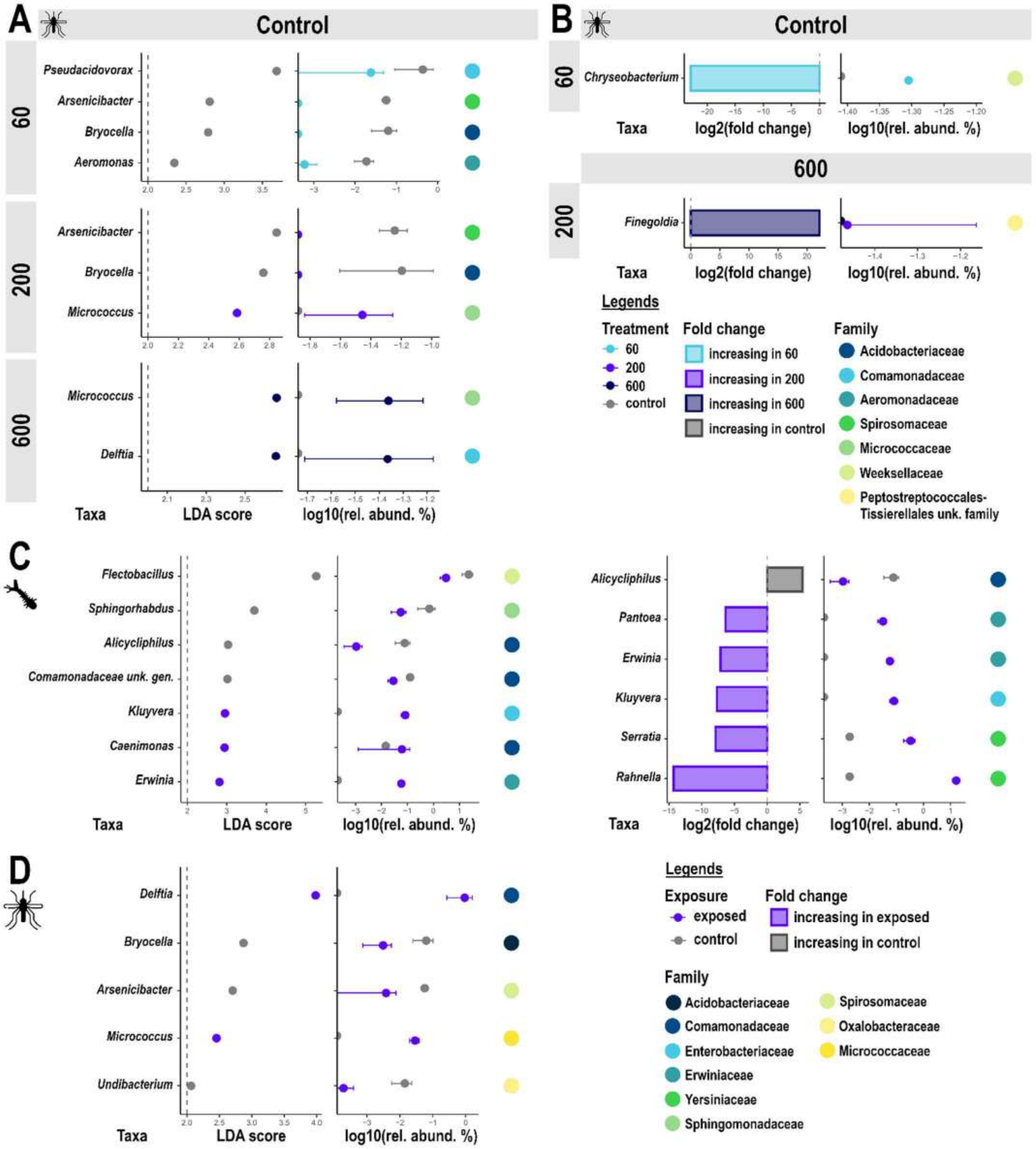

**Figure S8. Linear discriminant analyses for larvae and female microbiota. A.** Linear discriminant analyses performed for multiple group contrasts, including unexposed vs. individual concentration categories and pairwise comparisons between concentration categories, for the whole females. LDA scores are detailed alongside mean relative abundance per OTU and condition (as log10 values). For each OTU, we precise its taxonomic family. **B.** Differential abundance of genus-level OTUs between the unexposed group and each MP concentration category, for the whole females. We display only OTU with a p-value < 0.05 for the Wald test (including a Benjamini-Hochberg correction). Differential abundance, presented as log2(fold change), are detailed alongside mean relative abundance per OTU and condition (as log10 values). For each OTU, we precise its taxonomic family. **C.** LDA scores and differential abundance between unexposed and exposed groups, for the larvae. Metrics are detailed alongside mean relative abundance per OTU and condition (as log10 values). For each OTU, we precise its taxonomic family. **D.** LDA scores between unexposed and exposed groups, for the whole females. Differential abundance is not presented here, as no OTU exhibited a p-value < 0.05 for the Wald test.

### Supplementary figure 9

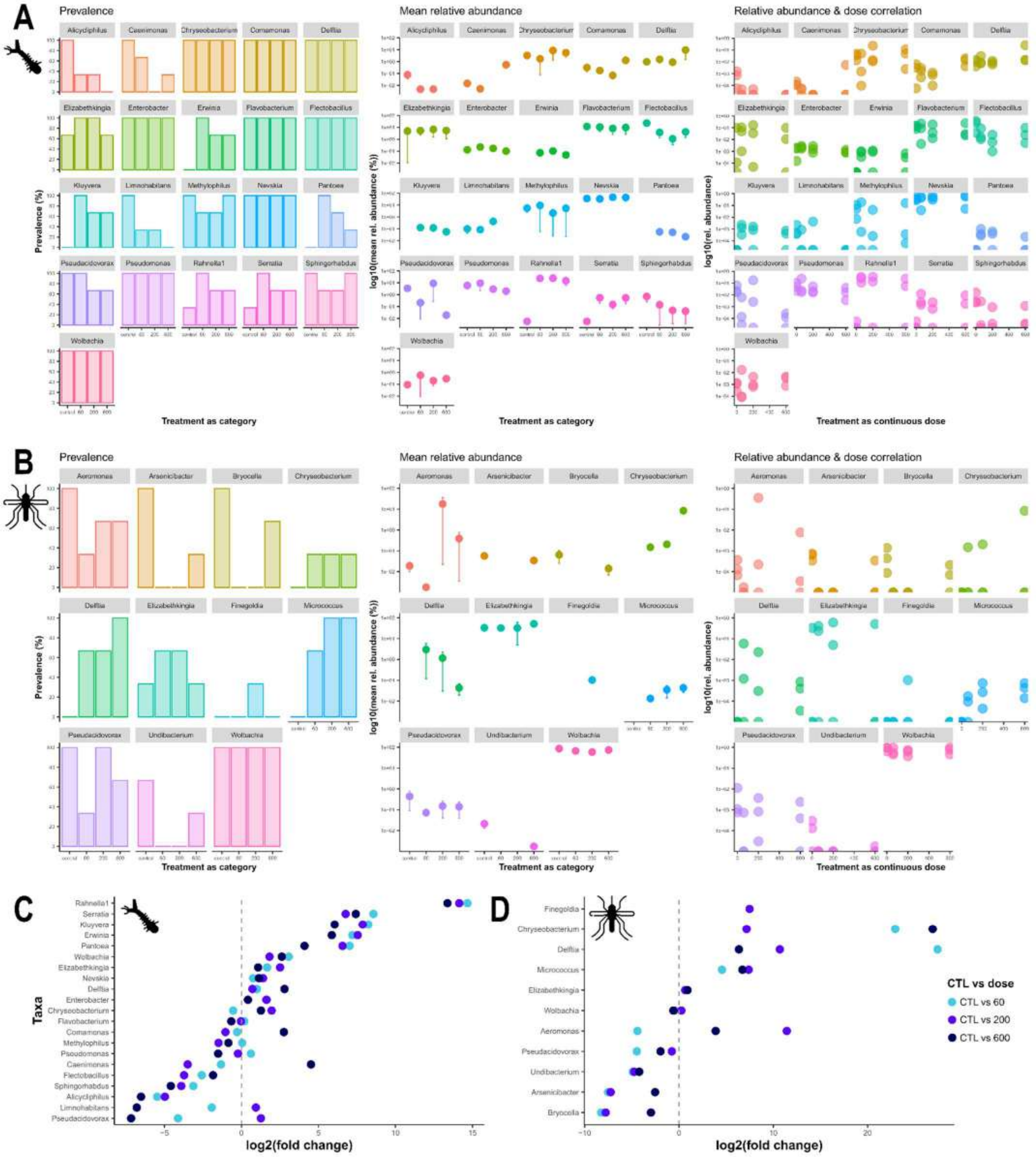

**Figure S9. Details about the prevalence and relative abundance of OTUs highlighted as significant by the linear discriminant analysis and differential abundance analysis, according to the MP concentrations**, for the larvae (A) and the whole females (B). Relative abundance is presented as mean and raw values according to MP concentration categories and doses, respectively. For each OTU, the differential abundance between the unexposed group and each MP concentration category, as a log2(fold change) value, is plotted for the larvae (C) and the whole females (D).

**Supplementary table 1**

**Table S1. Sequencing quality metrics for larvae and adult female samples across exposure conditions** (Control, PE60, PE200, PE600), including raw read counts, Q30 base-quality percentages, mapped read counts (with mapping rate), and GC content.

| Larvae | | | | | Females | | | | |
|---|---|---|---|---|---|---|---|---|---|
| **Sample** | **Raw_reads** | **Q30** | **Mapped_reads_M** | **GC_content** | **Sample** | **Raw_reads** | **Q30** | **Mapped_reads_M** | **GC_content** |
| **ControlL_rep1** | 64.61 M | 90.3 % | 46.35 M (71.7%) | 46.1 % | **ControlF_rep1** | 53.82 M | 93.4 % | 49.10 M (91.2%) | 29.4 % |
| **ControlL_rep2** | 45.73 M | 90.4 % | 35.10 M (76.8%) | 45.4 % | **ControlF_rep2** | 51.85 M | 92.8 % | 45.64 M (88.0%) | 30.1 % |
| **ControlL_rep3** | 28.80 M | 91.6 % | 23.96 M (83.2%) | 39.2 % | **ControlF_rep3** | 46.93 M | 94.7 % | 45.07 M (96.0%) | 24.8 % |
| **PE60L_rep1** | 44.38 M | 89.4 % | 34.74 M (78.3%) | 47.6 % | **PE60F_rep1** | 37.32 M | 93.0 % | 35.02 M (93.9%) | 26.3 % |
| **PE60L_rep2** | 56.53 M | 90.3 % | 49.71 M (87.9%) | 39.6 % | **PE60F_rep2** | 56.53 M | 92.8 % | 49.71 M (87.9%) | 30.2 % |
| **PE60L_rep3** | 44.74 M | 91.1 % | 34.46 M (82.2%) | 37.5 % | **PE60F_rep3** | 56.02 M | 94.0 % | 51.21 M (91.4%) | 29.4 % |
| **PE200L_rep1** | 53.63 M | 91.3 % | 41.92 M (78.2%) | 46.7 % | **PE200F_rep1** | 57.02 M | 94.7 % | 53.03 M (93.0%) | 26.1 % |
| **PE200L_rep2** | 41.10 M | 90.6 % | 32.80 M (79.8%) | 42.3 % | **PE200F_rep2** | 62.81 M | 92.6 % | 57.05 M (90.9%) | 29.0 % |
| **PE200L_rep3** | 44.04 M | 91.9 % | 36.81 M (83.6%) | 41.3 % | **PE200F_rep3** | 47.48 M | 94.3 % | 45.37 M (95.6%) | 25.2 % |
| **PE600L_rep1** | 45.44 M | 91.4 % | 31.81 M (70.0%) | 45.7 % | **PE600F_rep1** | 50.89 M | 92.4 % | 45.01 M (88.5%) | 30.7 % |
| **PE600L_rep2** | 37.21 M | 90.2 % | 30.56 M (82.1%) | 42.2 % | **PE600F_rep2** | 74.30 M | 93.3 % | 69.05 M (93.0%) | 27.5 % |
| **PE600L_rep3** | 38.94 M | 90.3 % | 32.12 M (82.5%) | 38.9 % | **PE600F_rep3** | 53.38 M | 93.7 % | 49.19 M (92.2%) | 27.6 % |

**Supplementary table 2**

**Table S2. Differentially Expressed Genes (DEGs) and their log2 Fold Change (FC) values.**

Table S2.1. Up- and down-regulated genes after exposure to 60 MP/mL in larvae

| VectorBase ID | Log2FC | Gene name | Description/activity | Regulation |
|---|---|---|---|---|
| CQUJHB006533 | 6.09 | *ncRNA gene* | - | Up-regulated |
| CQUJHB017497 | -5.36 | *Lysozyme c-1* | Lysozyme activity | Down-regulated |
| CQUJHB009037 | -5.24 | *Cecropin-A* | Antibacterial peptide | Down-regulated |
| CQUJHB009018 | -4.37 | *Defensin-C* | Antimicrobial peptide | Down-regulated |
| CQUJHB010736 | -3.38 | *CD109 antigen* | Endopeptidase inhibitor activity | Down-regulated |

Table S2.2. Up- and down-regulated genes after exposure to 200 MP/mL in larvae

| VectorBase ID | Log2FC | Gene name | Description/activity | Regulation |
|---|---|---|---|---|
| CQUJHB006533 | 5.58 | *ncRNA gene* | - | Up-regulated |
| CQUJHB016570 | 4.70 | *Uncharacterized LOC6043750* | Unknown | Up-regulated |
| CQUJHB014569 | 1.36 | *General odorant-binding protein 56d-like* | Odorant-binding protein | Up-regulated |
| CQUJHB017497 | -5.17 | *Lysozyme c-1* | Lysozyme activity | Down-regulated |
| CQUJHB004825 | -3.57 | *Bumetanide-sensitive sodium-(potassium) chloride cotransporter* | Transmembrane transporter activity | Down-regulated |
| CQUJHB001196 | -3.02 | *ncRNA gene* | - | Down-regulated |
| CQUJHB004192 | -2.48 | *Actin-87E* | Structural constituent of cytoskeleton (actin filament) | Down-regulated |
| CQUJHB009710 | -1.39 | *Endochitinase transcript variant X2* | Glycosyl hydrolase (chitinase) activity | Down-regulated |

Table S2.3. Up- and down-regulated genes after exposure to 600 MP/mL in larvae

| Gene ID | Log2FC | Gene name | Description/activity | Regulation |
|---|---|---|---|---|
| CQUJHB006533 | 6.89 | *ncRNA gene* | - | Up-regulated |
| CQUJHB009037 | -5.01 | *Cecropin-A* | Antibacterial peptide | Down-regulated |
| CQUJHB004857 | -4.56 | *Ejaculatory bulb-specific protein 3* | Small soluble chemosensory protein | Down-regulated |
| CQUJHB009018 | -3.91 | *Defensin-C* | Antimicrobial peptide | Down-regulated |
| CQUJHB017497 | -3.63 | *Lysozyme c-1* | Lysozyme activity | Down-regulated |
| CQUJHB012233 | -3.04 | *Uncharacterized LOC6036550* | Potential role in protein-protein interactions and signal transduction | Down-regulated |
| CQUJHB005028 | -2.60 | *Ficolin-1* | Pattern recognition receptor (lectin involved in innate immunity) | Down-regulated |

Table S2.4. Up- and down-regulated genes after exposure to 60 MP/mL in females

| VectorBase ID | Log2FC | Gene name | Description/activity | Regulation |
|---|---|---|---|---|
| CQUJHB009037 | 2.24 | *Cecropin-A* | Antibacterial peptide | Up-regulated |
| CQUJHB015711 | 3.22 | *Struthiocalcin-1* | EF-hand calcium-binding protein | Up-regulated |
| CQUJHB016620 | 5.44 | *Uncharacterized LOC6045020* | Protein present in salivary glands with unknown function | Up-regulated |
| CQUJHB016664 | 9.20 | *Perlucin* | C-type lectin (calcium-dependent carbohydrate-binding protein) | Up-regulated |
| CQUJHB013841 | -4.22 | *Trypsin 5G1* | Serine-type endopeptidase activity | Down-regulated |
| CQUJHB005980 | -3.09 | *Serine protease SP24D* | Serine-type endopeptidase activity | Down-regulated |

Table S2.5. Up- and down-regulated genes after exposure to 200 MP/mL in females

| VectorBase ID | Log2FC | Gene name | Description/activity | Regulation |
|---|---|---|---|---|
| CQUJHB016664 | 9.487 | *Perlucin* | C-type lectin (calcium-dependent carbohydrate-binding protein) | Up-regulated |
| CQUJHB016620 | 9.032 | *Uncharacterized LOC6045020* | Protein present in salivary glands with unknown function | Up-regulated |
| CQUJHB011416 | 3.142 | *Proline-rich protein 2-like* | Metalloendopeptidase activity | Up-regulated |
| CQUJHB015711 | 2.336 | *Struthiocalcin-1* | EF-hand calcium-binding protein | Up-regulated |
| CQUJHB005549 | 2.349 | *Trans-1,2-dihydrobenzene-1,2-diol dehydrogenase* | Oxidoreductase (dehydrogenase) activity | Up-regulated |
| CQUJHB005571 | 1.333 | *Macoilin, transcript variant X2* | Neuronal signal transduction | Up-regulated |
| CQUJHB006281 | -1.785 | *Peptide transporter family 1* | Transmembrane transporter activity | Down-regulated |
| CQUJHB014960 | -1.809 | *Uncharacterized LOC119766992* | Potentially involved in ERAD and ER–Golgi trafficking | Down-regulated |
| CQUJHB008375 | -2.088 | *Aminopeptidase N* | Metalloendopeptidase activity | Down-regulated |
| CQUJHB010407 | -2.427 | *ncRNA gene (lncRNA)* | - | Down-regulated |
| CQUJHB006747 | -2.715 | *Histone H1.3* | Linker histone (chromatin compaction & DNA binding) | Down-regulated |
| CQUJHB007612 | -3.139 | *Putative vitellogenin receptor* | Endocytic receptor for vitellogenin (LDL receptor family – yolk protein uptake) | Down-regulated |
| CQUJHB014381 | -3.418 | *Trypsin-7* | Serine-type endopeptidase activity | Down-regulated |
| CQUJHB007996 | -19.820 | *Uncharacterized LOC119770630, transcript variant X1* | Unknown | Down-regulated |

Table S2.6. Up- and down-regulated genes after exposure to 600 MP/mL in females

| VectorBase ID | Log2FC | Gene name | Description/activity | Regulation |
|---|---|---|---|---|
| CQUJHB016664 | 3.47 | *Perlucin* | C-type lectin (calcium-dependent carbohydrate-binding protein) | Up-regulated |
| CQUJHB010486 | -1.11 | *Uncharacterized LOC119770236* | Unknown | Down-regulated |

**Supplementary table 3**

**Table S3. Detailed information on the total number of raw and quality-filtered sequences, as well as the mean number of sequences and ASVs and OTUs per sample type.**

| Sample type | Raw sequence counts | Quality-passed sequences | Percentage of quality-passed sequences | Mean number of sequences per sample ± SE | Total number of ASVs | Mean number of ASVs per sample ± SE (min - max) | Total number of genus-level OTUs | Mean number of genus-level OTUs per sample ± SE (min - max) | Total number of samples |
|---|---|---|---|---|---|---|---|---|---|
| Larvae | 691,522 | 312,969 | 45.26 | 45,387 ± 6,600 | 864 | 240 ± 40 (82 - 418) | 61 | 35.6 ± 2.3 (25 - 49) | 12 |
| Whole females | 824,408 | 767,529 | 93.10 | 58,220 ± 5,874 | 550 | 121 ± 25 (57 - 380) | 83 | 27.8 ± 2.7 (18 - 47) | 12 |
| Female midguts | 2,584,439 | 2,243,608 | 86.81 | 54,759 ± 3,198 | 1,342 | 146 ± 18 (58 - 492) | 337 | 33.4 ± 2.3 (8 - 69) | 38 |

**Supplementary table 4**

**Table S4. Details regarding statistical tests performed on relative abundance and prevalence of Wolbachia and Elizabethkingia OTUs**, for the larvae, the whole females, and female midguts, separately, according to the MP exposure status and the MP concentration levels. NA: not applicable.

| | | | | Wilcoxon-Mann-Whitney | | Spearman's rank correlation | | Fisher's test |
|---|---|---|---|---|---|---|---|---|
| Sample type | OTU | Response variable | Explanatory variable | W | p-value | rho | p-value | p-value |
| **Larvae** | Wolbachia | Abundance relative | Exposure | 12 | 0.864 | - | - | - |
| | Wolbachia | Abundance relative | Concentration (doses) | - | - | 0.281 | 0.3768 | - |
| | Wolbachia | Prevalence | Exposure | - | - | - | - | NA |
| **Larvae** | Elizabethkingia | Abundance relative | Exposure | 6 | 0.711 | - | - | - |
| | Elizabethkingia | Abundance relative | Concentration (doses) | - | - | 0.113 | 0.757 | - |
| | Elizabethkingia | Prevalence | Exposure | - | - | - | - | 0.455 |
| **Whole females** | Wolbachia | Abundance relative | Exposure | 19 | 0.373 | - | - | - |
| | Wolbachia | Abundance relative | Concentration (doses) | - | - | -0.151 | 0.639 | - |
| | Wolbachia | Prevalence | Exposure | - | - | - | - | NA |
| **Whole females** | Elizabethkingia | Abundance relative | Exposure | 2 | 1 | - | - | - |
| | Elizabethkingia | Abundance relative | Concentration (doses) | - | - | 0.353 | 0.493 | - |
| | Elizabethkingia | Prevalence | Exposure | - | - | - | - | 1 |
